# Human hippocampal CA3 uses specific functional connectivity rules for efficient associative memory

**DOI:** 10.1101/2024.05.02.592169

**Authors:** Jake F. Watson, Victor Vargas-Barroso, Rebecca J. Morse-Mora, Andrea Navas-Olive, Mojtaba R. Tavakoli, Johann G. Danzl, Matthias Tomschik, Karl Rössler, Peter Jonas

**Author notes:** Correspondence to either J.F.W. or P.J.

## Abstract

The human brain has remarkable computational power. It generates sophisticated behavioral sequences, stores engrams over an individual’s lifetime, and produces higher cognitive functions up to the level of consciousness. However, so little of our neuroscience knowledge covers the human brain, and it remains unknown whether this organ is truly unique, or is a scaled version of the extensively studied rodent brain. To address this fundamental question, we determined the cellular, synaptic, and connectivity rules of the hippocampal CA3 recurrent circuit using multicellular patch clamp-recording. This circuit is the largest autoassociative network in the brain, and plays a key role in memory and higher-order computations such as pattern separation and pattern completion. We demonstrate that human hippocampal CA3 employs sparse connectivity, in stark contrast to neocortical recurrent networks. Connectivity sparsifies from rodents to humans, providing a circuit architecture that maximizes associational power. Unitary synaptic events at human CA3–CA3 synapses showed both distinct species-specific and circuit-dependent properties, with high reliability, unique amplitude precision, and long integration times. We also identify differential scaling rules between hippocampal pathways from rodents to humans, with a moderate increase in the convergence of CA3 inputs per cell, but a marked increase in human mossy fiber innervation. Anatomically guided full-scale modeling suggests that the human brain’s sparse connectivity, expanded neuronal number, and reliable synaptic signaling combine to enhance the associative memory storage capacity of CA3. Together, our results reveal unique rules of connectivity and synaptic signaling in the human hippocampus, demonstrating the absolute necessity of human brain research and beginning to unravel the remarkable performance of our autoassociative memory circuits.

## Introduction

We all experience the remarkable performance of the human brain. It generates exquisite behavioral sequences, stores memories over a lifetime, and provides us with our complex cognitive functions such as imagination and consciousness. Yet how cells and circuits generate these incredible phenomena remains unknown. In particular, we do not know whether the human brain is simply a scaled version of the extensively studied rodent brain (DeFelipe, 2011; Herculano-Houzel et al., 2015), or whether its uniqueness is produced by the specific properties of cells (Eyal et al., 2016), dendrites (Gidon et al., 2020; Mertens et al., 2024), or synapses (Campagnola et al., 2022; Molnár et al., 2016, 2008; Testa-Silva et al., 2014). Distinguishing between these possibilities requires structural and functional analysis of living human brain tissue. However, only very few experimental studies have directly examined the properties of human circuits. This is particularly striking for the hippocampus, which is the most intensively studied region of the rodent brain, yet of its cellular function in humans we know very little. The hippocampal CA3 region is the largest autoassociative network in the brain, with a fundamental role in memory storage (Lisman, 1999; Treves and Rolls, 1994). While its cellular, synaptic, and microcircuit properties were extensively studied in rodents (Guzman et al., 2016; Nakashiba et al., 2008), functional data from human CA3 are unavailable.

Currently, the only way to tackle these fundamental questions is to obtain direct recordings from living brain tissue extracted from human patients. It is generally believed that such measurements are only possible in neocortical tissue, which is thought to be unaffected by the underlying disease and better preserved during surgery (Campagnola et al., 2022; Peng et al., 2019; Planert et al., 2023; Seeman et al., 2018). However, recent work shows that hippocampal tissue extracted from patients during epilepsy surgery is heterogeneous, including patients in which the hippocampus is highly sclerotic, but also subjects in which the hippocampus appears largely unaffected (Roessler et al., 2016). Both pre-surgery magnetic resonance imaging (MRI) and post-surgery histological analysis show no differences between tissue in these patients and unaffected humans (Roessler et al., 2016). Such non-sclerotic samples provide a unique opportunity to determine the synaptic basis of memory storage and higher-order computations in the human hippocampus.

## Results

### Recording from neurons in the human hippocampus

To characterize cells, synapses, and circuits in the human CA3 region, we obtained hippocampal tissue blocks from 14 pharmaco-resistant temporal lobe epilepsy (TLE) patients who underwent unilateral temporal lobe resection surgery (**Fig. 1**; (Roessler et al., 2016)). This procedure resulted in the excision of neocortical and hippocampal tissue, which is immediately transferred to the laboratory for acute slice preparation and multicellular patch clamp-based functional circuit mapping (Guzman et al., 2016; Jiang et al., 2015; Peng et al., 2019; Perin et al., 2011; Seeman et al., 2018). By combining this analysis with post-hoc immunohistochemistry, visualization of recorded neurons, and expansion-based superresolution microscopy, we performed a detailed structural and functional characterization of the CA3 recurrent network in the human brain (**Fig. 1A–D**).

**Fig. 1.**
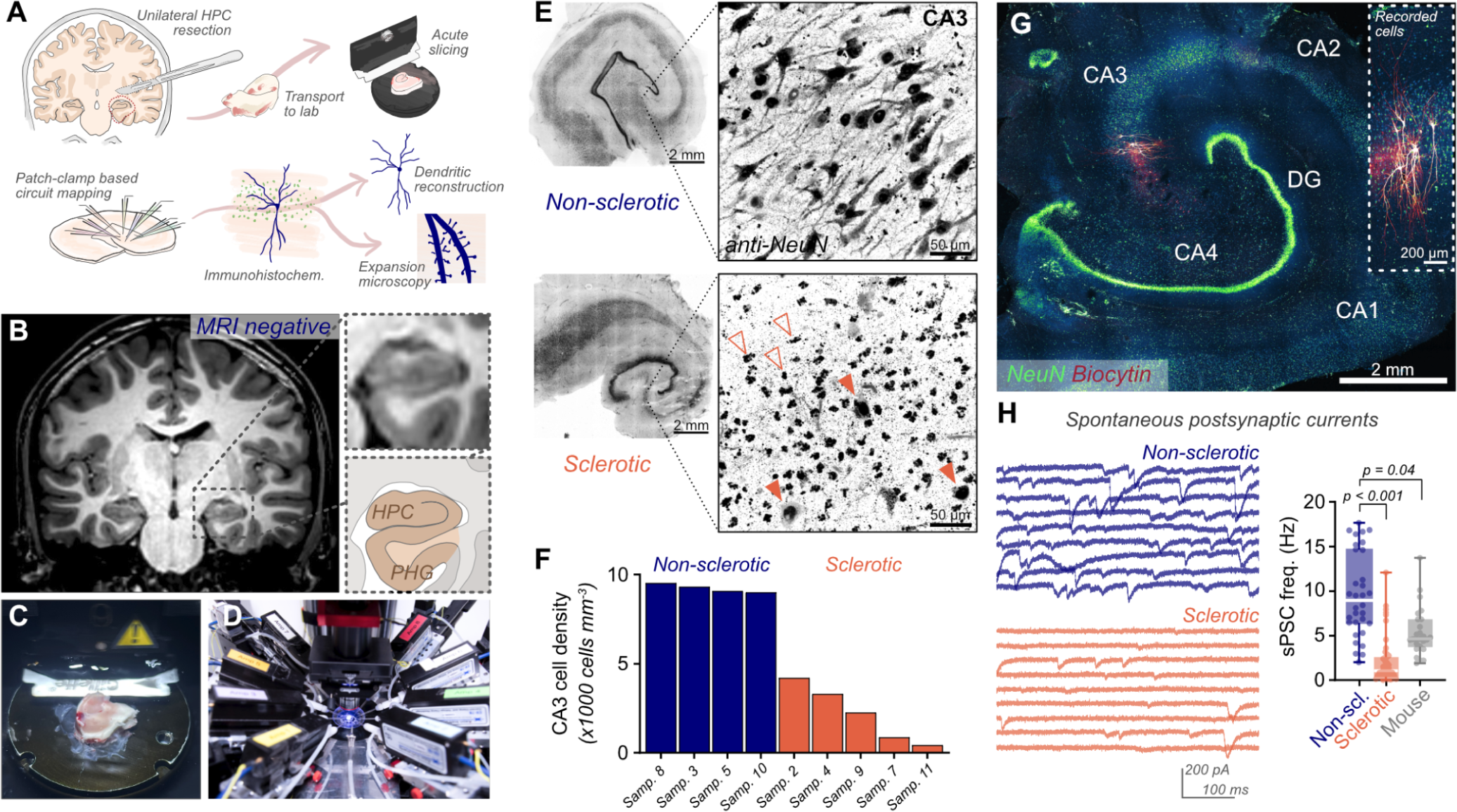
Using human hippocampal tissue from temporal lobe resection. **A**, Schematic of experimental procedure for analysis of human hippocampal circuits. **B**, Pre-surgery MRI brain image brain depicting tissue to be excised (schematic colored; HPC - hippocampus, PHG - parahippocampal gyrus). **C**, Hippocampal tissue block during sectioning to acute slices. **D**, Multicellular patch-clamp recording configuration. **E**, Anti-NeuN staining of acute human hippocampal slices shows (**F**) substantial and variable reductions in CA3 neuronal density in sclerotic tissue (orange), while non-sclerotic samples (blue) have a stable CA3 cell density. NeuN-positive neuronal remains are observed in sclerotic tissue, showing the extent of disease-induced cell loss (empty arrowheads, degenerated neurons; filled arrowheads, putatively healthy neurons). **G**, Example non-sclerotic recorded slice stained with anti-NeuN antibodies (blue-green LUT), and AF647-conjugated streptavidin to visualize biocytin-filled recorded cells (red; inset, magnified view). **H**, The rate of sPSCs at CA3 PNs is greatly reduced in sclerotic tissue when compared to non-sclerotic tissue, while human CA3 PNs (non-sclerotic) show a higher event rate than mouse CA3 PNs (Left, example traces of voltage-clamp recordings from human CA3 PNs. Mean ± SEM: non-sclerotic human: 9.6 ± 0.8 Hz, n = 32 cells; sclerotic human: 2.2 ± 0.4 Hz, n = 46 cells; mouse: 5.4 ± 0.5 Hz, n = 26 cells; Kruskal-Wallis test, p < 0.0001).

Pre-surgery MRI analysis, infrared differential interference contrast (IR-DIC) imaging during slice recording, and post-hoc immunohistochemical quantification of neuronal densities demonstrated that the population of patients was highly heterogeneous. In 8 out of 14 patients, hippocampal tissue showed sclerosis, with size reductions visible before surgery (‘MRI-positive’) and neuronal loss seen in tissue samples across the CA subfields including CA3 (**Fig. 1E**; (Blümcke et al., 2013)). In contrast, in 6 of 14 patients (‘MRI-negative’), no signs of hippocampal sclerosis were evident (**Fig. 1B, E**). Clinical histology analysis classed all these samples as lacking hippocampal sclerosis, and reported equivalent hippocampal cell densities and tissue structure to unaffected humans (**Supplementary Table 1**). We quantified cell density in CA3, and confirmed a high and stable density level (**Fig. 1F, G**; (West and Gundersen, 1990)). Furthermore, the frequency of spontaneous postsynaptic currents (sPSCs) in recorded CA3 pyramidal neurons (PNs) in this ‘non-sclerotic’ tissue was an order of magnitude higher than in sclerotic samples, as well as significantly higher than in mouse CA3 PNs, suggesting functionally preserved circuit wiring (**Fig. 1H**). Taken together, these analyses showed no evidence of disease induced damage to the hippocampal circuitry, corroborating the suggestion that CA3 is intact. This allowed us, for the first time, to examine the structure and function of human hippocampal microcircuits in the most physiological setting possible.

### Sparse synaptic connectivity in human CA3

To probe the functional synaptic connectivity in the human CA3, we used multicellular patch clamp-based circuit mapping. In total, we obtained 75 multicellular recordings from human hippocampal slices (10 octuples, 18 septuples, 15 sextuples, 7 quintuples, 6 quadruples, 12 triples, and 7 pairs across 14 patients, **Fig. 2A–C**), focusing our analysis only on recordings from the confirmed non-sclerotic patients (**Supplementary Table 1**). From these recordings, 204 neurons were confirmed to be CA3 PNs by electrophysiological and morphological criteria (**Supplementary Fig. 1**). In total, we found 10 putative monosynaptic connections in 1060 tested CA3 PN pairs, corresponding to a measured connection probability of 0.94% (**Fig. 2C**). To account for the effects of tissue slicing and to better approximate CA3 connectivity in the intact circuit, we performed a detailed morphological analysis of all recorded neurons and considered only neurons with extensive axonal staining as possible presynaptic partners (> 0.5 mm reconstructable axon, **Supplementary Fig. 2**; see also (Campagnola et al., 2022; Planert et al., 2023)). With this refinement, the corrected connection probability was 1.47% (10 out of 679 tested connections). Intersomatic distance between synaptically connected neurons varied over a wide range, with no evidence for distance dependence (**Supplementary Fig. 3**B), in contrast to other brain circuits (Planert et al., 2023), but consistent with CA3 measurements in rodents (Guzman et al., 2016). These results demonstrate that the human CA3 is a sparse but broadly connected recurrent network.

**Fig. 2.**
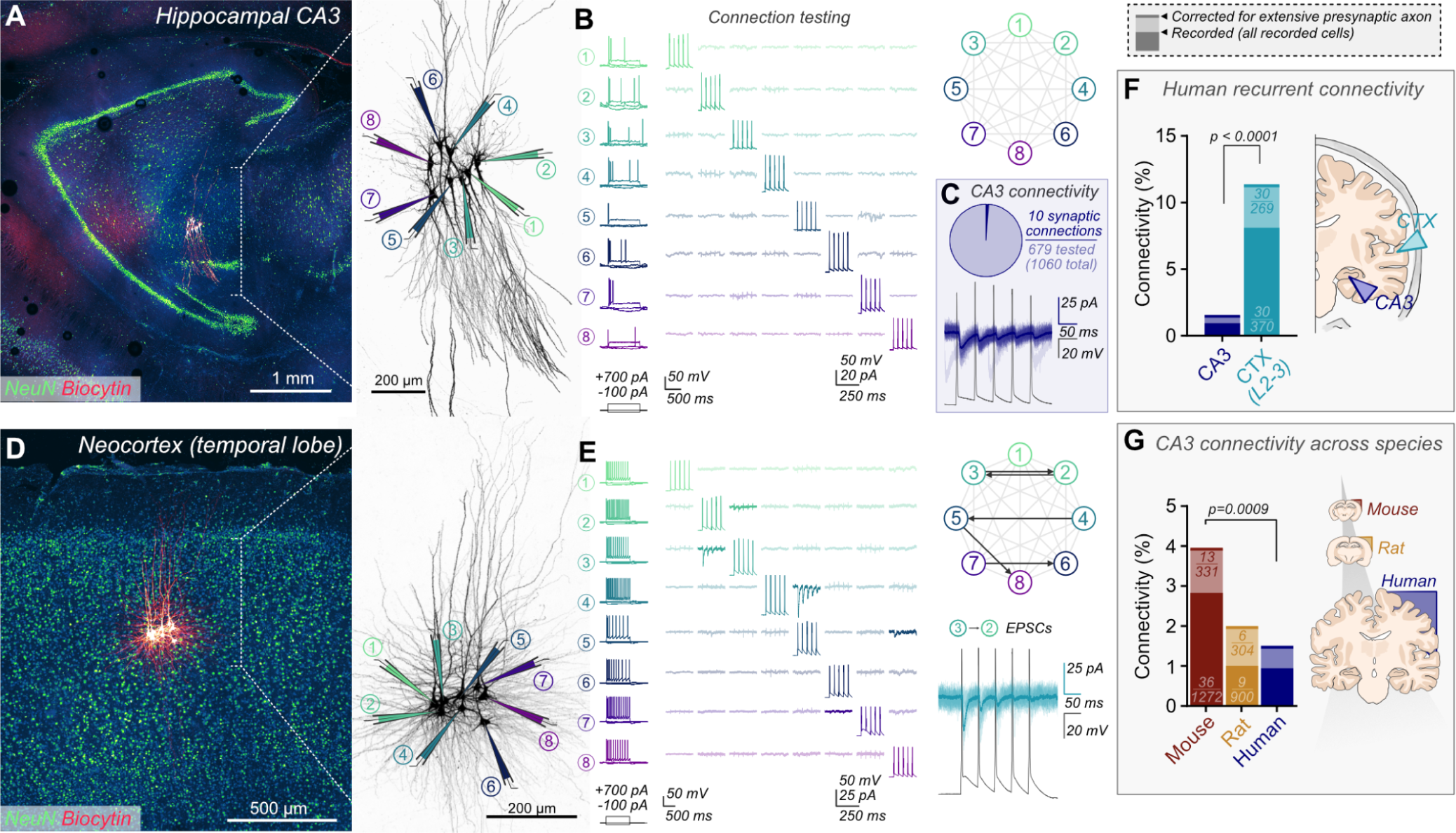
Sparse synaptic connectivity in human CA3. **A**, Human acute slice with octuple recording from CA3 (anti-NeuN, green; streptavidin-biocytin staining, red), with magnified view of recorded CA3 PNs (right) and annotated recording pipettes. **B**, Firing profile (left) and connectivity testing (right) for cells recorded in **A**. Averaged current traces in response to presynaptic cell spiking (diagonal) are shown. **C**, CA3 PN connectivity is low in human tissue (10/1060 tested connections, 10/679 connections after axon correction: 0.5 mm visible axon required for inclusion as a presynaptically tested neuron). Example of synaptic connection between human CA3 PNs is depicted. Presynaptic cell spiking (gray) is followed by reliable postsynaptic currents (individual traces are overlaid with the average response). **D**, Octuple recorded human neocortical neurons (layer 2-3), with corresponding firing properties and connection testing (**E**). Five synaptic connections are observed (bold traces), with connectivity scheme (upper) and magnified example synapse (lower) depicted right. **F**, Human neocortical L2-3 PN connectivity is an order of magnitude higher than between CA3 PNs (recorded: 30/370 tested connections, corrected: 30/269). Fisher’s exact test for both recorded and corrected connectivities are p < 0.0001. **G**, Sparse connectivity is observed in CA3 of both rodents and humans, with decreasing connectivity correlating with increasing brain size (mouse: 36/1272 tested connections, 13/331 axon corrected. Rat: 9/900 tested connections, 6/304 axon corrected. Mouse vs human Fisher’s exact test: p = 0.0009, total connectivity; p = 0.022, corrected connectivity).

Previous studies observed high connectivity between neocortical layer 2-3 PNs in humans (10–15%; (Campagnola et al., 2022; Kanari et al., 2024; Peng et al., 2024, 2019; Planert et al., 2023; Seeman et al., 2018)). To validate that sparse synaptic connectivity in human CA3 resulted from circuit architecture, rather than tissue or recording quality, we performed additional multicellular recordings from human layer 2-3 neocortical neurons under identical experimental conditions, using temporal lobe tissue resected during hippocampal surgery (**Fig. 2D, E**). We found dense recurrent connectivity in the human neocortex (corrected connectivity 11.3%, 30/269 tested connections, data from 2 patients), confirming previous observations (Campagnola et al., 2022; Peng et al., 2019; Planert et al., 2023), but in striking contrast to our findings in the human hippocampal CA3 region (p < 0.0001; **Fig. 2F**). Thus, sparse connectivity is a specific property of the human hippocampal CA3 circuit, whereas dense connectivity is characteristic for the human neocortex.

Sparse CA3 synaptic connectivity was previously demonstrated, in both rats (Guzman et al., 2016) and guinea-pigs (Miles and Wong, 1986). However, surprisingly high CA3 connectivity has recently been reported in mice (Sammons et al., 2024). To quantitatively analyze synaptic connectivity across species, we compared connection probability among humans, rats, and mice (**Fig. 2G, Supplementary Fig. 4**). When measured under identical experimental conditions, connectivity was highest in the mouse (measured: 2.83%; corrected: 3.93%), intermediate in the rat (measured: 1.00%; corrected: 1.97%), and lowest in humans (measured: 0.94%; corrected: 1.47%; **Fig. 2G**). Consistent with these findings, the branching density of CA3 axons was lower in humans than rodents (**Supplementary Fig. 2**). Therefore, not only does human CA3 employ much sparser connectivity than neocortical recurrent networks, but circuit architecture appears to sparsify across species with increasing brain size. Neocortical connectivity levels are comparable if not denser in humans than mice (Campagnola et al., 2022; Seeman et al., 2018), and formation of dense interconnectivity has been considered a mechanism for ‘uniquely human networks’ (Kanari et al., 2024). Our results suggest that the CA3 recurrent circuit uses a different mechanism to support human cognition.

### Reliable and precise synaptic transmission in human CA3

Next, we examined the functional properties of cells and synapses in the human CA3 (**Fig. 3**; **Supplementary Fig. 5, 6**). As our current understanding of hippocampal function is almost entirely based on analysis of rodent tissue, we first characterized the basic features of human CA3 PNs. Passive membrane properties of CA3 PNs showed similarities between humans and rodents. Human CA3 neurons showed greater cell capacitance, lower input resistance, and higher current threshold of action potential (AP) initiation (**Supplementary Fig. 5, 6B**). However, the membrane time constant was similar, indicating that differences were generated by larger surface area rather than differences in specific membrane properties. Active membrane properties were more distinct. Human CA3 PNs showed broader single APs, more pronounced adaptation, and narrower interspike interval distributions during repetitive firing (**Supplementary Fig. 6**). Thus, spiking of CA3 PNs showed a higher degree of temporal precision in humans than in rodents.

**Fig. 3.**
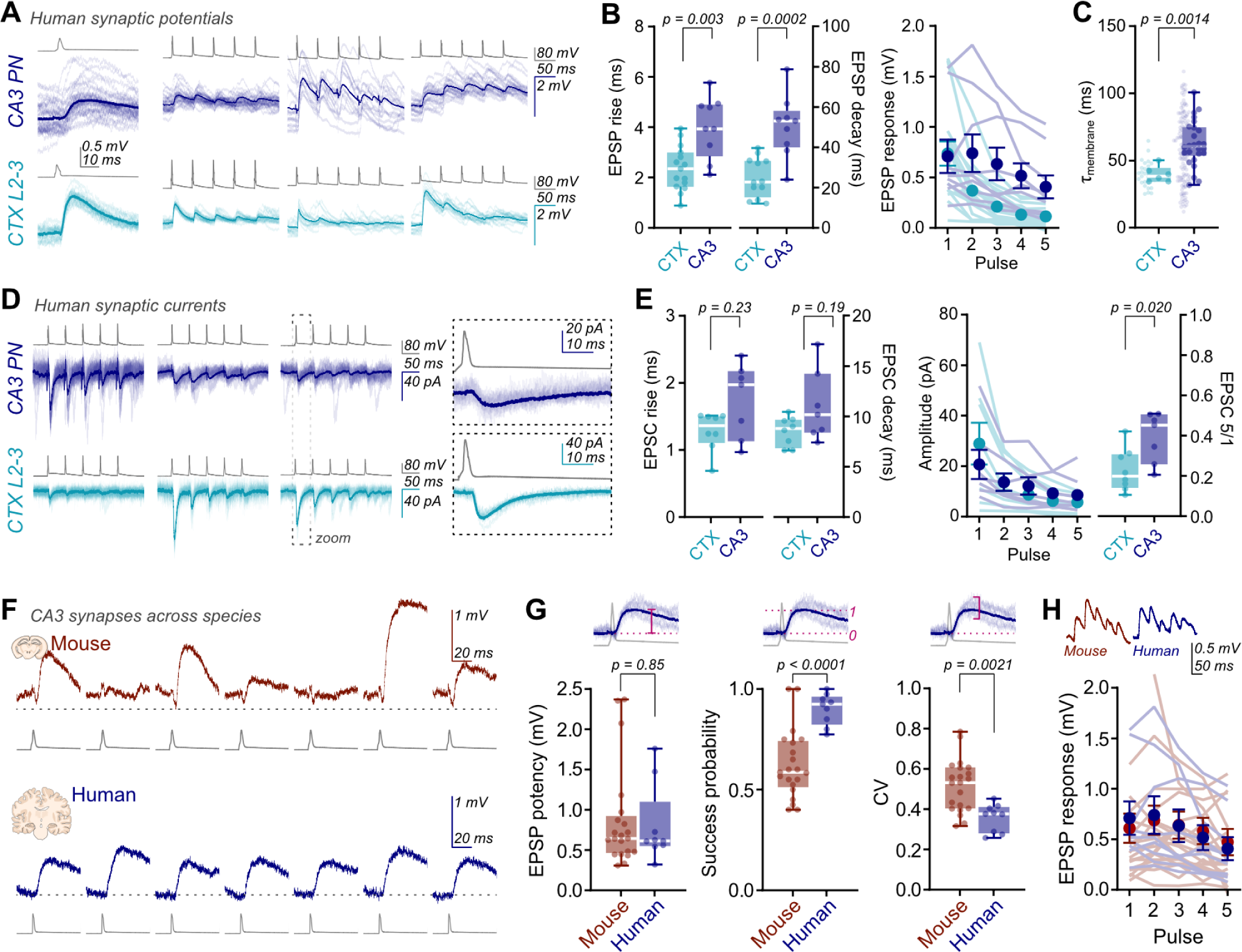
Reliable and precise synaptic transmission in human CA3. **A,** Unitary connections between human CA3 or neocortical layer 2-3 neurons (traces show individual sweeps overlain with average response), show (**B**) slower time course in human CA3 than human neocortical layer 2-3 (CTX; left, EPSP 20–80% rise time - CTX: 2.4 ± 0.2 ms, n = 15; CA3: 4.0 ± 0.4 ms, n = 9; Mann-Whitney test, p = 0.003. Right, EPSP decay time constant - CTX: 25.1 ± 2.8 ms, n = 12; CA3: 50.6 ± 5.1 ms, n = 9; Mann-Whitney test, p = 0.0002). The resulting EPSP response to 20-Hz train stimulation is less depressing in CA3 (right, EPSP5/1 ratio - CTX: 0.16 ± 0.03, n = 15; CA3: 0.58 ± 0.12, n = 9; Mann-Whitney test, p<0.0001). **C**, EPSP kinetics reflect the slower membrane time constant of large, integrating CA3 PNs (average per slice presented, with all cells depicted in light color. CTX: 39.8 ± 2.5 ms, n = 6 slices, n = 30 cells; CA3: 62.4 ± 3.3 ms, n = 26 slices, n = 133 cells; Mann-Whitney test, p = 0.0014). **D**, Unitary EPSCs from human slices. **E**, EPSC kinetics are comparable between areas (left, EPSC 20–80% rise time - CTX: 1.3 ± 0.1 ms, n = 8; CA3: 1.7 ± 0.2 ms, n = 7; Mann-Whitney test, p = 0.23. EPSC decay time constant - CTX: 8.5 ± 0.5 ms, n = 8; CA3: 11.0 ± 1.3 ms, n = 7; Mann-Whitney test, p = 0.19), while CA3 synapses are less depressing than neocortical synapses (right, EPSC5/1 amplitude ratio - CTX: 0.23 ± 0.04, n = 8; CA3: 0.39 ± 0.05, n = 7; Mann-Whitney test, p = 0.020). **F**, Example sweeps from a CA3 synapse of mouse (upper, red) and human (lower, blue), which show (**G**) similar strength in both species (mouse: 0.87 ± 0.14 mV, n = 21; human: 0.80 ± 0.16 mV, n = 9; Mann-Whitney test, p = 0.82), yet human synapses are substantially more reliable, with minimal synaptic failures (probability of success - mouse: 0.62 ± 0.04, n = 21; human: 0.90 ± 0.03, n = 9; Mann-Whitney test, p = 0.0002), and less variation in the amplitude of successes (coefficient of variation, CV - mouse: 0.52 ± 0.03, n = 21; human: 0.36 ± 0.02, n = 9; Mann-Whitney test, p = 0.0021). **H**, Average responses to 20-Hz trains are similar between species.

Our recordings present the first examples of paired synaptic recordings from the human hippocampus (**Fig. 3**). In comparison to the well characterized neocortical synapses, excitatory postsynaptic potentials (EPSPs) evoked between recorded human CA3 PNs had notably slower rise and decay time (**Fig. 3B**). CA3 PNs showed a markedly slower membrane time constant than neocortical neurons (p = 0.0014; **Fig. 3C**), suggesting differences in postsynaptic membrane properties as a potential mechanism. To further distinguish between effects of synaptic conductance and postsynaptic membrane properties, we measured the kinetics of excitatory postsynaptic currents (EPSCs), and found far less kinetic difference than between EPSPs (EPSC rise and decay kinetics: p = 0.23 and 0.19, respectively; **Fig. 3D, E**). Finally, human CA3 synapses were less depressing than neocortical synapses during 20-Hz train presynaptic stimulation (p < 0.0001 for EPSPs and 0.02 for EPSCs; **Fig. 3B, E**). Together therefore, human CA3 PNs provide a broader window for temporal summation. Both the cellular and synaptic properties of CA3 PNs produce a recurrent circuit with greater integrative power in human hippocampal CA3 than in human neocortical layer 2-3.

To identify potentially unique properties of synaptic signaling in humans, we analyzed unitary EPSPs and EPSCs in different species (**Fig. 3F–G**). Direct comparison of mouse and human CA3 synaptic properties showed similar EPSP and EPSC amplitudes and kinetics across species (**Fig. 3G; Supplementary Fig. 7**). However, there was a major difference in the reliability and amplitude precision of synaptic transmission. Human CA3–CA3 synapses were strikingly reliable, with a success probability of 0.9, while mouse synapses showed a significantly lower success probability (0.62; p < 0.0001; **Fig. 3G**). Although the synaptic potency (i.e. the amplitude of successes) was equivalent, the coefficient of variation (CV) of success amplitudes was significantly smaller in humans than in mice (0.36 vs. 0.52; p = 0.002). Both differences were corroborated in voltage-clamp recordings (**Supplementary Fig. 7**A–C). Analysis of reliability and amplitude precision of human neocortical synapses revealed that potency, success probability, and CV of successes were not significantly different to human CA3 (**Supplementary Fig. 8**). Taken together, human synapses show much lower synaptic fluctuations than mouse synapses, with both greater reliability and precision. Reliable and precise transmission appears to be a general feature of synaptic signaling in human recurrent networks (Hunt et al., 2023). Therefore, our recordings identify unique functional properties employed by the human brain.

### Scaling of recurrent connectivity from mice to humans

Which factors determine sparse synaptic connectivity in the human CA3? In a broad, random network (Erdős and Rényi, 1959), connectivity can be estimated anatomically as the number of inputs per cell (N_inputs_), divided by the total number of cells in the network (N_cells_). Thus, connectivity would be predicted to be proportional to N_inputs_ but inversely proportional to N_cells_. We tested these predictions in the CA3 network. Stereology analyses indicated that CA3 contains ∼110,000 CA3 PNs per hemisphere in mouse ((Attili et al., 2022); J.F.W, unpublished), ∼300,000 CA3 PNs per hemisphere in rat (Attili et al., 2022; Boss et al., 1987), and ∼1.8 M CA3 PNs per hemisphere in humans ((González-Arnay et al., 2024; Seress, 1988; Simić et al., 1997; West, 1993; West and Gundersen, 1990) see **Methods**). Therefore CA3 undergoes a massive evolutionary expansion in cell number ((Herculano-Houzel et al., 2015); see **Fig. 5A**).

We next attempted to directly measure N_inputs_ (**Fig. 4**). The size of the neuronal dendritic arbor is known to expand from rodents to humans (Benavides-Piccione et al., 2020; Wang et al., 2016), but the contribution of scaling to CA3 function is unknown. We analyzed the dendritic arbors of human CA3 neurons by confocal microscopy and reconstruction (**Fig. 4A–D**). Human neurons were highly heterogeneous, with examples of extreme basal dendrite extension or apparent ‘inverted’ topology (**Fig. 4B**). To determine the cable length contributing to the CA3 recurrent system, we considered dendrites covered by simple spines in *stratum oriens* and *stratum radiatum*, while excluding proximal dendrites decorated by thorny excrescences (mossy fiber input) and distal dendrites entering *stratum lacunosum-moleculare* (perforant path input). In a subset of CA3 PNs, we observed that dendrites crossed the granule cell (GC) layer (**Supplementary Fig. 1**), which we also presumed to receive perforant path input. We corrected for dendritic cutting by calculating the expected length of all dendrites terminating at the slice surface. Total dendritic length receiving CA3 input increased from 6.5 mm per PN in mice up to 15.3 mm in humans (**Fig. 4D**), therefore cell size expansion resulted in a 2.37±0.08-fold extension of CA3 recurrent dendrite per cell from mice to humans.

**Fig. 4.**
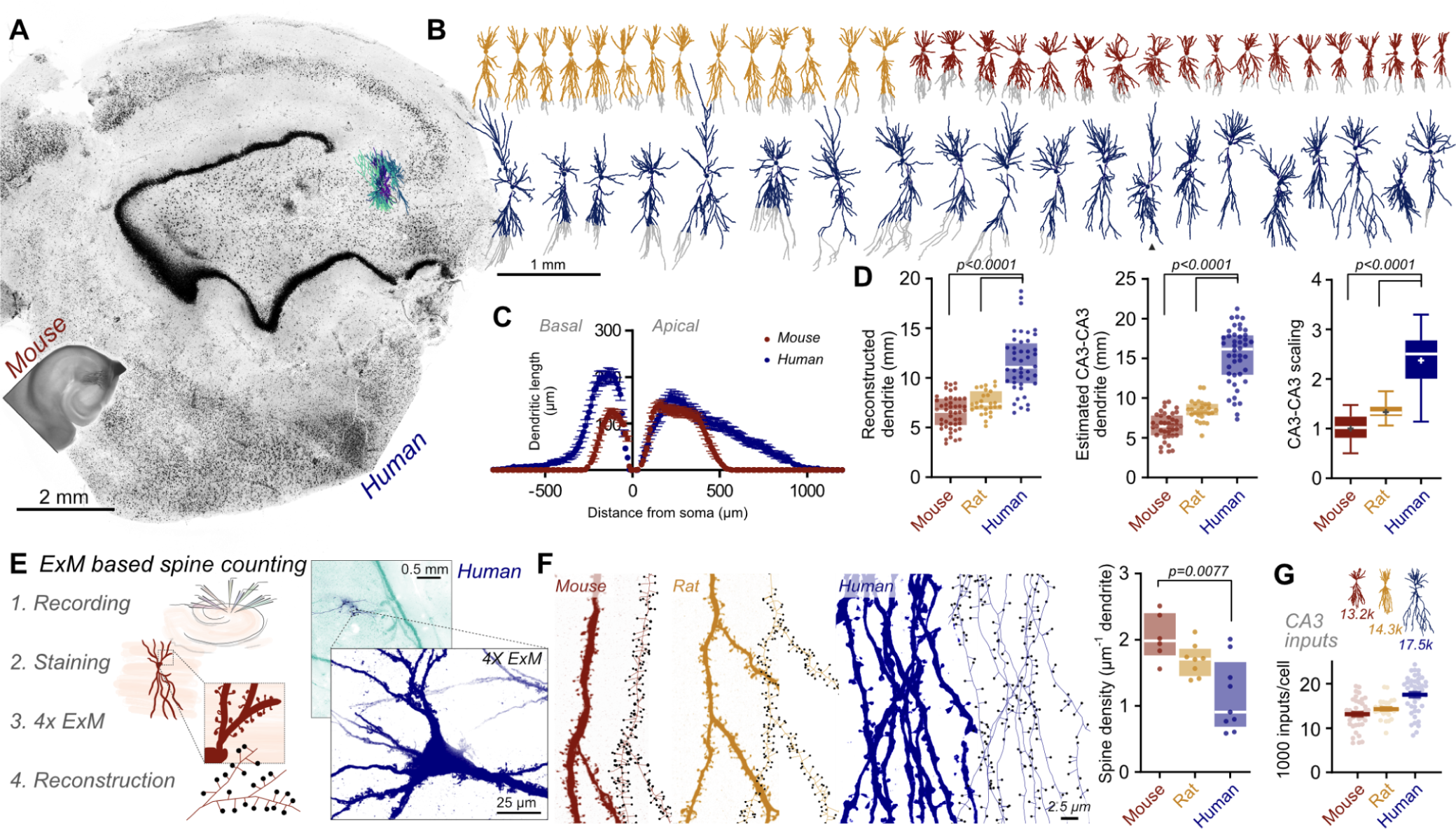
Human CA3 PNs have longer dendrites but lower spine density. **A**, Human and mouse hippocampal slice images demonstrate the expansion in hippocampal size in humans. NeuN-stained human tissue (confocal) is depicted in scale with a DAB-stained mouse slice. Reconstructed human neurons are depicted (colored). **B**, Array of reconstructed mouse (red), rat (yellow), and human (blue) CA3 PN skeletons, demonstrating cell size expansion across species. Somata are depicted as circles, while dendrites not likely to be contributing to the CA3 recurrent circuit are colored gray. A neuron with putative inverted topology (i.e. reversed direction of basal and apical dendrites) is indicated (triangle). **C**, Quantified dendritic length with respect to the soma (zero) for reconstructed mouse and human neurons shows an increase in the abundance and length of basal dendrites in human CA3 PNs, while predominantly an extension in length of human apical dendrites (mouse (red), n = 33; human (blue), n = 28 cells). **D**, The total dendritic length reconstructed from recorded human CA3 PNs is greater than that of rodent PNs(left, mean ± SEM: mouse, 6.5 ± 1.7 mm, n = 46 cells; rat, 7.5 ± 1.2 mm, n = 27 cells; human, 11.5 ± 3.0 mm, n = 44 cells; Kruskal-Wallis test, p < 0.0001), with a corresponding extension of dendritic length predicted to form the CA3 recurrent circuit (center: mouse, 6.5 ± 1.7 mm, n = 39 cells; rat, 8.5 ± 1.3 mm, n = 27 cells; human, 15.3 ± 3.6 mm, n = 44 cells; Kruskal-Wallis test, p < 0.0001). Species-scaled dendritic length (normalized to mouse), shows an approximately 2.5-fold increase in recorded dendrite in the CA3 recurrent system of human PNs over rat or mouse neurons (right, mean marked as +; mean ± SEM: mouse 1.00 ± 0.04, n = 39 cells; rat, 1.31 ± 0.04, n = 27 cells; human, 2.37 ± 0.08, n = 44 cells; Kruskal-Wallis test, p < 0.0001). **E**, 4x expansion microscopy (ExM) approach for analysis of spine density on recorded neurons (left), and (right) recorded human slice (anti-NeuN, green; biocytin-filled cells, blue) with ExM image of recorded human CA3 PN (indicated on overview), showing detailed morphology of dendritic spines and thorny excrescences. Scale bar indicates pre-expansion size. **F**, Imaged and reconstructed dendrites and spines (circles) from CA3 PN dendrites. Scale bar indicates pre-expansion specimen size. The density of dendritic spines on human CA3 dendrites is lower than mouse dendrites (mean ± SEM: mouse, 2.04 ± 0.14 spines µm^−1^, n = 6 cells; rat, 1.69 ± 0.09 spines µm^−1^, n = 8 cells; human, 1.14 ± 0.18 spines µm^−1^, n = 9 cells (3 patients); Kruskal-Wallis test, p = 0.0096). **G**, The estimated inputs per CA3 neuron (estimated CA3 dendrite length x mean spine density) increases only minimally from mice to humans (mouse: 13,200; rat: 14,300; human: 17,500 inputs per cell).

We then counted dendritic spines on the relevant dendritic branches (**Fig. 4E–G**). Individual spines are difficult to unequivocally determine with diffraction-limited light microscopy. Therefore, we optimized a 4x expansion microscopy (ExM) protocol (4x ProExM; (Tillberg et al., 2016)) that enabled superresolution analysis of dendritic architecture from recorded neurons (**Fig. 4E**). Using this protocol, biocytin-streptavidin labeled neurons in previously fixed and cleared human tissue could be expanded and imaged at 4x increased resolution even one year after initial sample mounting. We could visualize both thorny excrescences (mossy fiber input) and simple spines on primary dendrites of human CA3 neurons (recurrent collateral input; **Fig. 4E–F**). Simple spines occurred at similar densities on both basal and apical dendrites of rodent neurons (**Supplementary Fig. 9**), but unexpectedly, spine density was ∼2-fold lower on human than rodent CA3 PNs (**Fig. 4F**). This decreased spine density in human tissue parallels electron microscopy (EM) observations from neocortical PNs (Loomba et al., 2022). Therefore, lower excitatory input density appears to be a conserved feature of excitatory neurons across the human brain.

Based on these results, we estimated the average number of spines per complete CA3 PN across species, calculating 13,200 spines per CA3 PN in mice, 14,300 in rats, and 17,500 in humans (**Fig. 4G**). Thus, the evolutionary expansion of the hippocampus is accompanied by a massive increase in cell number, a substantial increase in dendritic length, and a decrease in spine density, overall leading to a moderate increase in the number of synaptic inputs. Taken together, these results indicate that connectivity in the CA3 circuit is not governed by a simple scaling rule, but rather by multiple scaling rules with different slopes and directions (**Fig. 5A**).

**Fig. 5.**
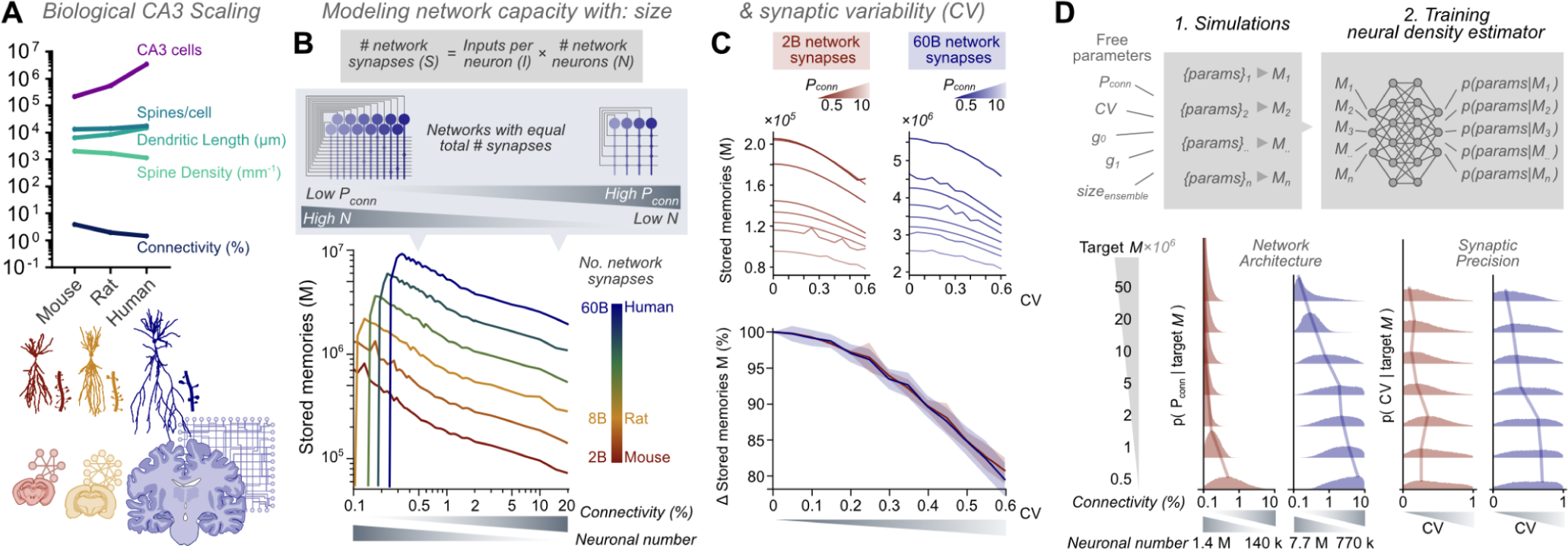
Sparse connectivity and reliable synaptic signaling increase memory storage capacity of a CA3 PN network model. **A**, Measured scaling of CA3 network parameters across species. **B**, Theoretical CA3 network models with fixed numbers of total synapses in the network (variable number of nodes and inputs per node) show maximal pattern storage and retrieval with lower connectivity (high N), than with higher connectivity. This applies to networks of all sizes, equivalent to biological CA3 networks across species (colored). **C**, Adding variability to synaptic transmission (increasing CV) reduces the memory storage capacity of networks of all sizes (upper). Networks (from **B**) with varying connectivities (shades) show equivalent effects. This reduction occurs by an equivalent proportion, as seen from normalized data (lower). Lines and shading represent mean and SD of normalized capacity for networks with different structure of equivalent size (colored) **D**, Simulation-based inference (SBI) optimisation of CA3 network properties to achieve increasing maximal memory capacities show peak shifts of optimal parameter distributions to lower connectivity (high N, low P_conn_) and low CV with increasing memory capacity requirements, supporting **B** and **C**. Lines follow maxima of distributions. Networks with 2 billion (red) and 60 billion (blue) synapses were tested.

Using these numbers, we can predict synaptic connectivity for a theoretical CA3 network (**Supplementary Fig. 10**A). Assuming that the number of spines equates the number of inputs (Bourne et al., 2013; Rigby et al., 2023; Sorra and Harris, 1993) and that ipsilateral and contralateral inputs contribute equally to synaptic connectivity (Blackstad, 1956; Qiu et al., 2024), we predict an average connection probability of 5.99% for mouse, 2.65% for rat, and 0.52% for human (**Supplementary Fig. 10**A; see **Methods** for caveats and limitations). These values approximate experimental measurements surprisingly well, in line with the view of CA3 as a broadly connected network, and supported by axon morphology across species (Ishizuka et al., 1990; Kondo et al., 2009; Witter, 2007; Wittner et al., 2007). Interestingly, connectivity levels akin to the neocortex are only possible with much more focal connectivity. As a human CA3 PN has only 17.5 k inputs, 15% local connectivity would imply that one neuron receives all its input from within 3% of the total human CA3 (113 k neurons;**Supplementary Fig. 10**B), while 1.5% local connectivity (as measured) broadens this influence to 33% of the network. Therefore, the sparse connectivity we identify in human hippocampal CA3 reflects the mutually exclusive circuit architectures between CA3 and neocortex: sparse and broad connectivity in CA3 versus dense and ‘columnar’ connectivity in the neocortex (Mountcastle, 1997; Douglas and Martin, 2004; Perin et al., 2011; Jouhanneau et al., 2018; Seeman et al., 2018; Campagnola et al., 2022; Planert et al., 2023). Such a network arrangement likely maximizes the associative power of the hippocampus, facilitating association between parallel, spatially separated information streams.

### Theoretical determinants of maximal network capacity

Our results reveal key characteristics of human CA3 circuits: sparse connectivity and reliable synaptic signaling. We sought to understand the influence of these properties on memory storage capacity, using an established theoretical model of CA3 memory function (**Fig. 5**; (Hopfield, 1982; Bennett et al., 1994; Guzman et al., 2016)). In this Hopfield-type network, patterns were stored by Hebbian synaptic plasticity, and retrieved from incomplete or degraded versions by pattern completion. Evolutionary expansion of CA3 could have adopted alternative routes: increasing connectivity and consequently input number per neuron, or preferring sparser connectivity (P_conn_) between greater neuronal numbers (N). To test the efficiency of these options for memory storage, we analyzed networks with a constant total number of synapses, changing connectivity and cell number reciprocally (**Fig. 5B**). Thus, we tested the theoretical pattern storage capacity of small, densely connected networks (high P_conn_, low N) up to large, sparsely connected networks (low P_conn_, high N) (**Fig. 5B**). This approach allows more legitimate comparison of different network structures, surpassing comparisons between networks with equivalent N but differing P_conn_, which will have vastly different synapse numbers available. Capacity of the network was measured as the maximal number of patterns that could be stored without interfering with reliable recall of previous patterns. Recurrent networks of all sizes, from 2 billion synapses (approximating mouse CA3), to 60 billion synapses (approximating human CA3) consistently showed greater storage capacity for lower connectivity combined with higher cell number than for higher connectivity combined with lower cell number (**Fig. 5B**), except at unrealistically low connectivity values, in which networks were silent due to insufficient recurrent activity (**Fig. 5B**, bottom, left). Therefore, for a theoretical CA3 network, an evolutionary drive for improved memory capacity would prioritize expansion of neuronal number over an increase in network interconnectivity. We extended this analysis to examine the effect of synaptic reliability on pattern storage. Increasing synaptic variability (CV) reduced the memory storage capacity for networks of all sizes in a proportional manner (**Fig. 5C**). Together, our results suggest that maximal storage capacity in a theoretical recurrent network is achieved by sparse connectivity, increasing neuronal number rather than each neuron’s input number, and enhancing synaptic reliability, the properties we experimentally recorded in human CA3.

To test whether our conclusions hold more generally, we performed a further analysis avoiding any need to constrain model parameters (P_conn_, CV, ensemble size, threshold (g_0_) and inhibition (g_1_); **Fig. 5D**). This ‘simulation-based inference’ approach (SBI; (Papamakarios and Murray, 2016; Gonçalves et al., 2020)) is based on Bayesian inference, and uses model simulations with arbitrary parameter values to train a neural density estimator, which can subsequently compute the distribution of parameters most likely to reach a given memory capacity value M. Using this unbiased approach, when increasing the target value of M, we observed consistent shifts in the peak of parameter distributions to smaller values of both connectivity and CV, in both small and large networks (**Fig. 5D**). Therefore achieving maximal network capacity is most likely with high neuronal numbers, sparse connectivity, and reliable synaptic transmission. These results corroborate the idea that the human CA3 follows an efficient coding design, maximizing associational power and memory storage capacity.

### Pathway-specific expansion of CA3 synaptic inputs

While the CA3 recurrent connectivity is proposed to store representations of our experiences, this information is thought to be relayed from dentate gyrus GCs (Rolls, 2013, 2018). Input from GCs reaches CA3 via the powerful synapses of the hippocampal mossy fibers (Vandael and Jonas, 2024). While some signaling properties of human mossy fiber synapses have been determined (Pelkey et al., 2023), the rules of their connectivity in humans are unknown, and will critically influence CA3 function. As with CA3 PNs, GC numbers show substantial evolutionary expansion, from ∼460,000 GCs per hemisphere in mice (Amrein et al., 2004; Ihunwo and Schliebs, 2010) to ∼15 M per hemisphere in humans (Simić et al., 1997; West, 1993; West and Gundersen, 1990). Compared to the well-known rodent hippocampal architecture, the human GC layer has greatly extended in length, maintaining a thin layer with a high GC density. In contrast, the CA3 layer has fanned out along the deep-superficial axis (**Fig. 1G, 6A, B**). The clear separation between mossy fiber tract (*stratum lucidum*) and cell body layer (*stratum pyramidale)* observed in mice has been lost in human CA3, with PN somata and mossy fibers intermingled ((González-Arnay et al., 2024; Lim et al., 1997); **Fig. 6B**). As a result, we observed thorny excrescences on both apical and basal dendrites of human CA3 PNs, implying that mossy fiber input occurs on a wider range of subcellular domains of postsynaptic target cells (**Fig. 6A, B**; see also (Lauer and Senitz, 2006; Lu et al., 2013)).

**Fig. 6.**
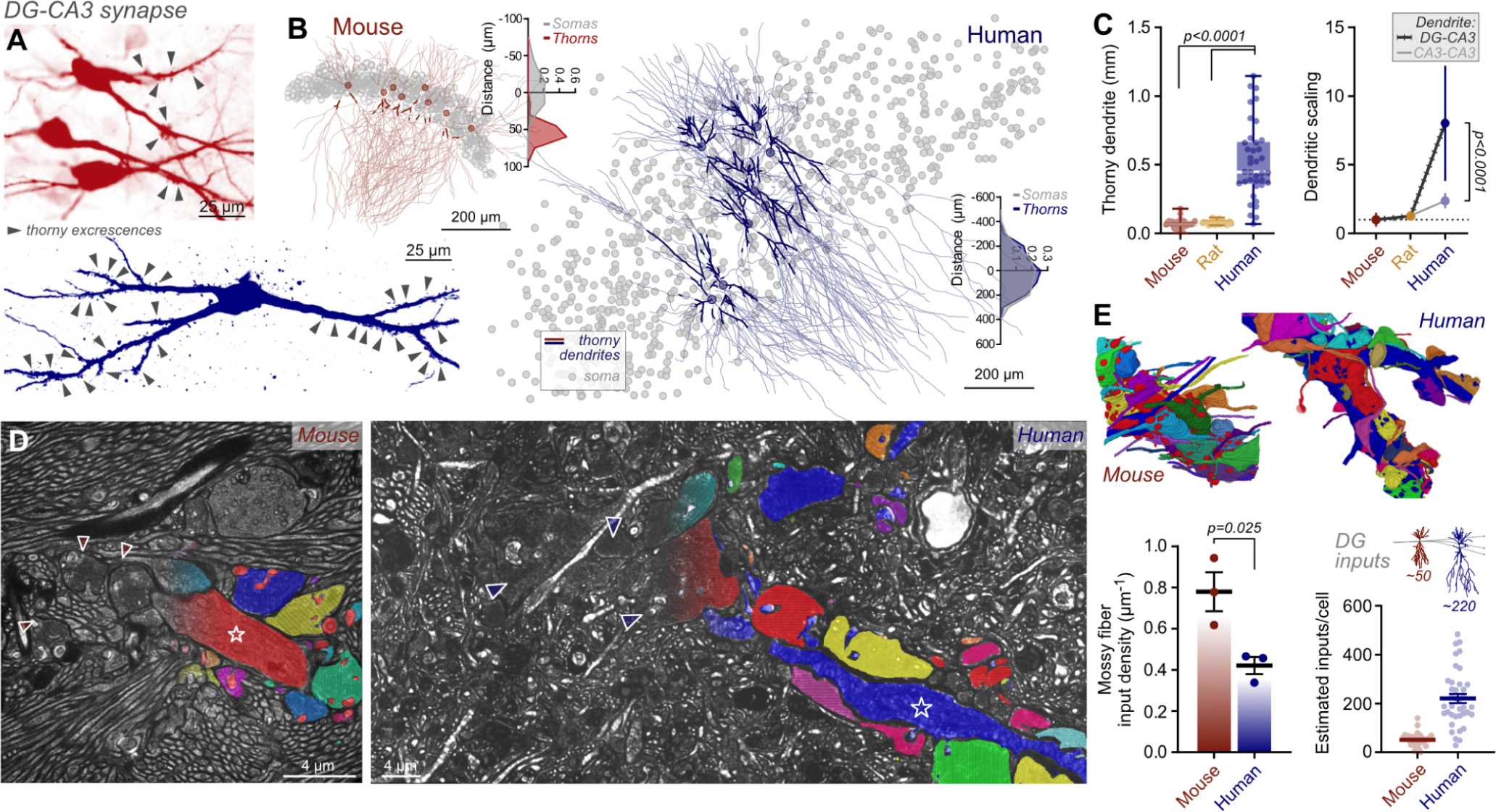
Expanded mossy fiber-CA3 input in the human brain. **A**, Both mouse (upper) and human (lower) CA3 PNs display complex spines (‘thorny excrescences’, arrowheads) on primary apical dendrites of biocytin-labeled recorded cells. **B**, Left, thorny dendrites of mouse neurons (bold lines on recorded cell reconstructions) segregate from the pyramidal cell layer (somata – gray circles). Right, human thorny dendrites overlap with the expanded *stratum pyramidale* and extend throughout the somatic layer. Histograms depict distributions of somas (gray) and thorny dendrite (colored). **C**, Length of reconstructed thorn-covered dendrite is expanded in humans compared to mouse CA3 PNs (mouse: 0.065 ± 0.005 mm, n = 39; human: 0.52 ± 0.04 mm, n = 38), equating to a 8-fold expansion in dendritic coding space (left – absolute values; right – normalized to mouse mean length; human dendritic scaling: thorny dendrite, 8.0 ± 0.7, CA3 dendrite, 2.4 ± 0.1, Mann-Whitney test, p < 0.0001). **D**, LICONN-based tissue imaging allows reconstruction of thorny dendrite (starred) and contacting mossy fiber boutons (indicated with triangles or colored segmentation, scale bar reflects pre-expansion length). Segmentation is included on half of images for presentation of raw data. **E**, While bouton density appears lower in human tissue (boutons per unit length of thorny dendrite, mouse: 0.78 ± 0.09 μm^−1^, n = 3 cells; human: 0.42 ± 0.09 μm^−1^, n = 3 cells, unpaired t-test, p = 0.025), estimated GC inputs per CA3 PN increases vastly in the human brain (mouse: 51 ± 4; human: 221 ± 19).

The length of thorny dendrite on reconstructed PNs increased 8.0±0.7-fold from mice to humans, supra-proportional to cell size and the 2.4±0.1-fold increase for the CA3 recurrent collateral system (**Fig. 6C**). To confirm that thorny dendrites were indeed the target of mossy inputs in human tissue, we applied LICONN (Tavakoli et al., 2024), an iterative hydrogel-expansion technology, enabling synapse-level tissue reconstruction. An effective spatial resolution of <20 nm laterally (∼16-fold increased over confocal imaging) with high-fidelity tissue preservation and comprehensive structural labeling (pan-protein), allowed us to reconstruct the mossy fiber connectome of PN dendrites, which displayed extensive large boutons contacting thorny excrescences in both mouse and human (**Fig. 6D**). Bouton density on human PNs was reduced to 54% of densities on mouse PNs (0.42 vs 0.78 boutons per µm dendritic length in humans vs. mice; p = 0.025; **Fig. 6E**). While this is a further example of lower excitatory input density in human tissue, this effect is eclipsed by the massive expansion in thorny dendritic length. We calculated that a human CA3 PN receives on average ∼220 GC mossy fiber inputs, in comparison to just ∼50 for a mouse CA3 PN (see also (Amaral et al., 1990); **Fig. 6E**). At the level of GC-CA3 connectivity, these values correspond to 0.011% and 0.0015% of GCs connecting to each CA3 PN in mice and humans respectively, suggesting connectivity is sparser at both GC and CA3 recurrent inputs to the human CA3 area. Individually however, human CA3 PNs experience a much greater level of GC innervation to those of the mouse brain. As a proposed key function of CA3 is association of incoming dentate gyrus input streams (Rolls, 2018), this striking expansion of human mossy fiber convergence has the potential to dramatically increase the associational power of human CA3.

## Discussion

Until now, the functional properties of cells, synapses, and circuits in the human hippocampus have not been characterized. It is often assumed that the mechanisms of synaptic signaling and information processing are conserved across species, and that investigating basic mechanisms in model organisms will provide insights into higher cognitive functions in humans. However, a proof for this assumption is lacking. Here, we have tackled this question and present functional microcircuit properties of the human hippocampal CA3 region, the largest autoassociative network in the brain, which plays a key role in learning, memory, and higher-order computations. To the best of our knowledge, this study reports the first recordings from synaptically connected pairs of neurons in the human hippocampus, revealing specific features of these connections that could only be observed through direct analysis of human brains. Our results were obtained from structurally intact tissue, providing the closest possible approximation of the physiological human brain. However, tissue inevitably comes from patients with disease background, which must be considered when interpreting the results.

We demonstrate that connectivity of human CA3 cannot be predicted from the rodent brain by a simple scaling rule. CA3 connectivity does not scale similarly to neocortical circuits, instead decreasing from mice to humans. In addition, discrete CA3 microcircuit properties change following diverse scaling relations, acting differently on neuron numbers, dendritic architecture, and input density. Notably, we find that the different hippocampal inputs to CA3 PNs do not expand uniformly, with distinct changes to recurrent collaterals and mossy fiber inputs.

The decreased excitatory input density between mice and humans that we observe at two hippocampal synapses parallels similar observations in the neocortex ((Loomba et al., 2022) **Fig. 4F, g**), suggesting it may be a wider principle of human PN architecture. Our results also reveal that the functional properties of unitary synaptic transmission in humans are distinct from those in rodents. Human CA3–CA3 recurrent synapses show highly reliable transmission, with a high proportion of successes and a small CV of success amplitude. This is different from rodents, where transmission at these synapses is unreliable (Allen and Stevens, 1994). Higher synaptic reliability has been also suggested in human neocortical layer 2-3 synapses (Hunt et al., 2023), and therefore may also be a wider principle of human synaptic transmission. Whether this reliability is related to either broader APs (**Supplementary Fig. 6**) or lower spine density in human circuits remains to be addressed.

Our results provide evidence that the specific properties of the human CA3 network have major advantages, increasing the computational power of the network. Classical models of the hippocampal CA3 region suggested that incoming information is transferred by hippocampal mossy fiber synapses, which induce association and storage of information in CA3–CA3 synapses by Hebbian synaptic plasticity (Marr, 1971; Rolls, 2018). Our results indicate that specialization of both synapses in the human brain may be critical for network performance. The extension of the mossy fiber termination zone may allow single cells to respond to a much larger number of synaptic input combinations, increasing single-cell computation. As a consequence, this could greatly enhance the ability of the network to perform pattern separation computations and to participate in combinatorial encoding. The human hippocampus has followed an efficient coding route of expanding neuronal number. Employing sparse but broad CA3–CA3 connectivity will improve pattern completion and allow the system to associate engrams in a distributed, hippocampus-wide manner. This is fundamentally different from the neocortex, in which dense connectivity within cortical modules or columns can enable fast and efficient local computations (Kanari et al., 2024). Together, we reveal distinct features of microcircuit function in the human hippocampus. Model organism research provides the backbone to understand our brain’s function, but direct measurements from human brain tissue are essential to reveal the full picture.

## Methods

### Human tissue samples and slicing

Human TLE tissue samples were obtained with informed patient consent, and approved by the Ethics Committee of the Medical University Vienna (MUW). A subsection of the resected tissue was sent to the neuropathological laboratory (Division of Neuropathology and Neurochemistry, Department of Neurology, Medical University of Vienna, MUW) for histological assessment, and sub-classification according to international league against epilepsy (ILAE) sclerosis standards (Blümcke et al., 2013)(see also **Supplementary Table 1**). MRI image is presented (**Fig. 1**) with informed consent from the patient and the Department of Biomedical Imaging and Image-guided Therapy, Division of Neuroradiology and Musculoskeletal Radiology (MUW).

Tissue blocks of human hippocampus and cortex were removed from the brain, and transported to the laboratory in high-sucrose artificial cerebrospinal fluid (aCSF, containing 64 mM NaCl, 25 mM NaHCO_3_, 2.5 mM KCl, 1.25 mM NaH_2_PO_4_, 10 mM D-glucose, 120 mM sucrose, 7 mM MgCl_2_, and 0.5 mM CaCl; osmolarity ∼334 mOsm) equilibrated with carbogen. 350-µm-thick acute slices were cut, before recovery at 35°C in high-sucrose aCSF for at least 30 mins. After recovery, slices were maintained in high-sucrose aCSF at room temperature (RT; 20–22°C) until recording. Cortical tissue blocks were cut similarly, with macroscopic trimming to best preserve the dendritic architecture of cortical pyramidal neurons.

### Animal tissue samples and slicing

All procedures were performed in strict accordance with institutional, national, and European guidelines for animal experimentation, approved by the Bundesministerium für Wissenschaft, Forschung und Wirtschaft of Austria. Wild-type C57BL/6J mice (RRID:IMSR_JAX:000664) and wild-type Wistar rats (RRID:RGD_13508588) of both sexes were used at postnatal (P) day 20–30 unless otherwise stated. Animals were sacrificed by decapitation under isofluorane anesthesia and brains were extracted rapidly in ice-cold high-sucrose aCSF equilibrated with carbogen. 350-µm-thick acute slices were cut using the ‘magic-cut’ quasi-transverse preparation (Bischofberger et al., 2006) in partially frozen high-sucrose aCSF, and recovered in high-sucrose aCSF at 35°C for 30–45 mins before maintenance in this solution at RT until recording.

### Electrophysiology

Slices were transferred to the recording chamber, and continuously perfused under gravity flow with recording aCSF (containing 125 mM NaCl, 25 mM NaHCO_3_, 2.5 mM KCl, 1.25 mM NaH_2_PO_4_, 25 mM D-glucose, 2 mM CaCl_2_, and 1 mM MgCl_2_, osmolarity ∼317 mOsm) continuously bubbled with carbogen. Patch-clamp recording pipettes (open tip resistances of 2–6 MΩ) were pulled from thick-walled borosilicate glass tubing (Hilgenberg, 2 mm OD, 1 mm ID, 1807542), and filled with intracellular solution (containing 135 mM K gluconate, 20 mM KCl, 0.1 mM EGTA, 2 mM MgCl_2_, 2 mM Na_2_ATP, 0.3 mM NaGTP, and 10 mM HEPES, adjusted to pH 7.28 with KOH; osmolarity ∼302 mOsm, with 0.2% (w/v) biocytin). Micropipettes were positioned manually using eight Junior 20ZR micromanipulators (Luigs and Neumann), signals recorded using four Multiclamp 700B amplifiers (Molecular Devices), and digitized at 20 kHz with Power 1401 data acquisition interfaces (Cambridge Electronic Design). Protocols were generated and applied using Signal 6.0 software (CED). All recordings were made at RT.

Synaptic connectivity was tested by eliciting action potentials in each recorded neuron in current-clamp mode in turn, while recording responses from all other neurons in either current- or voltage-clamp configurations. APs were elicited by brief current injection (2–5 ms) at a minimal level to reliably evoke single. Connectivity was tested by 20 Hz stimulation repeated at least 40 times. Monosynaptic connections were identified by reliable representation of synaptic responses in the average trace, and short latency and minimal temporal jitter. After recordings, pipettes were slowly retracted to form outside-out patches to ensure retention of intracellular biocytin for post-hoc staining. Recorded traces were analyzed using Stimfit (version v0.15.8; (Guzman et al., 2014)) or custom Matlab scripts. All voltage-clamp recordings were performed with a holding potential of −70 mV. sPSCs were detected using a template based analysis (minidet.m, Biosig toolbox, http://biosig.sf.net/; see (Jonas et al., 1993; Clements and Bekkers, 1997)). Success probabilities refer to the first pulse of 20-Hz train stimulation. CV analysis of EPSPs and EPSCs was performed by measuring peak amplitude variation (SD/mean) from success sweeps. EPSP and EPSC ‘potency’ represent the amplitude of successful sweeps excluding failures. EPSP ‘response’ refers to the total depolarisation after successive stimuli of a train, including effects of temporal summation. Synaptic current rise times are between 20–80% of maximal response, while decay kinetics are time constant of a monoexponential fit.

### Immunohistochemistry

After recording, slices were fixed with 4% (w/v) paraformaldehyde (PFA) in 0.1 M phosphate buffer and washed in 0.1 M PB after 24 h to stop the fixation reaction. Slices were blocked and permeabilised by incubation with solution containing 5% normal goat serum and 0.4% Triton X-100 (Sigma-Aldrich) in 0.1 M PB and stained with Alexa Fluor (AF) 647-conjugated streptavidin (1:300 diluted from 2 mg ml^−1^ stock, Invitrogen S32357) overnight in blocking/permeabilising solution. DAPI (4’,6-diamidino-2-phenylindole, dilactate, 0.1 µg ml^−1^ final concentration in PB) labeling was performed before clearing in CUBIC solution (Tainaka et al., 2014) before mounting on glass slides in CUBIC solution surrounded by a ring of Mowiol (Mowiol 4-88, Carl Roth, 713,2; made up according to manufacturer protocol with Glycerol and Tris(hydroxymethyl)aminomethane) to prevent exposure of CUBIC solution to air.

Primary antibody staining was included prior to streptavidin addition by overnight incubation of primary antibodies in blocking/permeabilisation solution before overnight incubation with both secondary antibodies and conjugated streptavidin. Human slices were routinely stained with anti-NeuN (diluted 1:300 from stock; Rabbit anti-NeuN, Thermo Fisher Scientific PA5-37407, RRID:AB_2554049), visualized with AF488-conjugated anti-rabbit secondary antibody (1:300 from stock; Goat anti rabbit AF488, Thermo Fisher Scientific A11034, RRID:AB_2576217). Slides were imaged on an ANDOR Dragonfly microscope (Oxford Instruments) equipped with a Zyla sCMOS camera. Overview images were taken using a 10x air objective (Nikon MRD00105, CFI P-Apo 10x, NA 0.45), while cellular morphology was captured using a 20x water-immersion objective (Nikon MRD77200, CFI P-Apochromat 20x, NA 0.95). Higher resolution images for analysis of dendritic spines and thorny excrescences were taken using a 40x water-immersion objective (Nikon MRD77410, Apochromat LWD λS 40x, NA 1.15). Tiles were stitched using Imaris Stitcher software (Oxford Instruments). A subset of rodent samples were imaged using DAB staining, as described in Guzman et al. 2016. A subset of cells were recorded with somata outside this layer, closer to the GC layer, which are considered CA4. Zero connections (90 tested) were observed between CA4 cells. Neuronal densities were measured by blinded manual segmentation of NeuN-stained neurons from image stacks taken using the 40x water immersion objective in the CA3c region of human slices.

### Morphological analysis

Neurons were manually reconstructed using Neutube analysis software (Feng et al., 2015). SWC files were analyzed using the FIJI (Schindelin et al., 2012) SNT plugin, or using custom MATLAB scripts. Dendritic identity was classified into apical or basal dendrites with either simple spines, thorny excrescences or no structures by manual inspection. A subset of recorded cells across multiple patients was fully reconstructed for morphological analysis, however all recorded neurons were visualized to determine axon-corrected connectivity. Corrected connectivity considers only neurons with at least 0.5 mm of traceable axon included as possible presynaptic cells, while all recorded neurons were considered possible postsynaptic partners. Complete dendritic length was estimated by calculation of ‘cut length’ on a dendrite by dendrite basis for each reconstructed cell. For each dendritic node terminating at the slice surface, the dendritic path length to the soma was measured and ‘cut-length’ was calculated as the mean downstream dendritic length of all uncut dendrites beyond the equivalent somatic path length. Thorny dendritic data were not corrected for dendritic cutting due to their location close to the soma.

### Expansion microscopy for spine density measurement

PFA-fixed and AF647-streptavidin stained slices were expanded using a modified version of ProExM (Tillberg et al., 2016), applicable for fixed, CUBIC-cleared, and mounted samples, even over one year post-fixation. Briefly, samples were first anchored by overnight incubation (∼16 h) with 0.1 mg ml−1 Acryloyl-X, before incubation with gel monomer solution for 3–5 h at 4°C. Monomer solution contained 7% sodium acrylate, 20% acrylamide, 0.1% methylene bisacrylamide 0.05% ammonium persulfate (APS), and 0.05% tetramethylethylenediamine (TEMED) in Milli-Q water. Samples were subject to gelation at 37°C for 3 h in a moist chamber before being trimmed to the region of interest, and digested by proteinase K (New England Biolabs, 1:100 dilution to 8 units ml^−1^) overnight at 37°C. Digestion buffer was removed by repeated washing in PBS at RT while shaking for 3–4 h. Streptavidin signal was amplified by incubation with biotinylated anti-streptavidin antibody (1:200 dilution, Vector Laboratories, VECBA-0500, RRID:AB_2336221) before addition of further AF647 conjugated streptavidin (1:300 dilution). Slices were expanded by incubation in MilliQ water before imaging on an ANDOR Dragonfly spinning disk microscope as above. Expansion factors were calculated by comparison of pre and post expansion images and were measured to be 3.9 ± 0.1, 3.9 ± 0.1, and 3.8 ± 0.1 for mouse, rat and human respectively (custom code, (Tavakoli et al., 2024)). Spines were either detected using a machine learning algorithm in ilastik (Berg et al., 2019) and manually corrected using Imaris 9 (Bitplane), or manually segmented using Neutube.

### Caveats of anatomical connectivity predictions

While dendritic spine density provides a good approximation of synaptic input, several factors add uncertainty to exact quantifications. EM analysis of human neocortex demonstrated an increase in the proportion of excitatory connections onto PN dendritic shafts, from 0.7% (mice) to 12.0% (humans) (Loomba et al., 2022). It is possible that this also occurs in the hippocampus, leaving our input numbers underestimated by around 10%. In addition, complex innervation arrangements complicate a definitive determination of input number from spine density measurements. Connected neocortical neurons for example, have a mean of 1.6 synaptic contacts per connection (Holler et al., 2021), causing an overestimation of presynaptic neuronal input number from spine counts. While connectomic analysis of CA3 is almost non-existent, multi-contact synapses have been suggested (Guzman et al., 2016). In CA1, which receives CA3 PN axons, spines appear to overwhelmingly receive a single input (Bourne et al., 2013), and multi-synaptic boutons (Rigby et al., 2023) or axons (Sorra and Harris, 1993) rarely contact the same cell twice. Finally, a substantial proportion of synapses in the brain may be functionally silent (Kullmann, 2003; Vardalaki et al., 2022), therefore likely causing overestimations of functional connectivity from structural data. Together, spine density is likely to be a good approximation of the number of CA3 presynaptic neurons in the recurrent circuit, but is not a definitive measure of input number.

### High resolution expansion microscopy for synapse reconstruction (LICONN)

Mossy fiber connections in human and mouse samples were reconstructed using high-resolution expansion with LICONN technology (Tavakoli et al., 2024), with slight adaptations for the human specimens as detailed below, and manual cell shape annotation.

Three wild-type mice (age 8–10 weeks) were anesthetized, first by isoflurane inhalation, then by intraperitoneal ketamine/xylazine (80–100 mg kg^−1^ and 10 mg kg^−1^ of body weight respectively), combined with subcutaneous metamizol (200 mg kg^−1^). Animals were transcardially perfused with RT PBS for 2 min followed by 6 min with RT PFA-fixative solution. Brains were postfixed for ≤ 12 h at 4°C and sectioned coronally at 50 µm thickness using a vibratome. Sections were quenched in 100 mM glycine for 6–8 h at 4°C. Human acute tissue slices (350 µm) were fixed immediately after slicing in RT fixative solution before quenching in 100 mM glycine-containing PBS for 30 min. Further handling was according to (Tavakoli et al., 2024) with adaptations for human tissue. Images were acquired using an ANDOR Dragonfly spinning disk microscope with a 40x objective lens as described above. CA3 primary dendrites were identified by their thorn-studded morphology, and both dendrite and large boutons encasing thorny excrescences were 3D segmented manually using VAST Lite (v1.4.0, (Berger et al., 2018)). Reconstructed dendritic length covered with thorns was measured in Neutube and used to determine the bouton density per unit length. 3D visualizations were created in VAST Lite.

### Estimates of neuronal numbers

Estimates of neuronal numbers in rodents were used from the Hippocampome database and recent meta analyses (Attili et al., 2022; Wheeler et al., 2015). Human neuronal densities used the mean values of separate stereology studies (Seress, 1988; Simić et al., 1997; West, 1993; West and Gundersen, 1990). Neuronal counts are reported for combined CA3 and CA2 fields. The estimate of neuronal numbers in human CA3 alone were acquired by measurement of the relative CA3 and CA2 *cornu ammonis* layer lengths in the hippocampal body from a detailed field parcellation study (González-Arnay et al., 2024) to determine the scaling factor: CA3 ≈ 67% of CA3-2.

### Mathematical modeling of CA3 function

Pattern completion analysis was performed in an autoassociative memory network model described in previous works (Bennett et al., 1994; Gibson and Robinson, 1992; Guzman et al., 2016; Hopfield, 1982). To account for the differences between hippocampal sizes, we modeled the network using realistic sizes. Due to the large number of excitatory neurons of the human hippocampus (> 10^6^) and the computational power it would require, instead of a simulation-based approach (such in Guzman et al., 2016), the analytical approximation of the model was used (Bennett et al., 1994). To make the comparison between network sizes consistent with basic biological properties of the hippocampus, we fixed the total number of synapses of the network, and distributed them in small highly connected networks (small n, high c), or increasingly bigger and less connected networks (high n, small c). Total number of synapses was fixed to 2, 4, 8, 16, 32, and 60·10^9^. Tested connectivity ranged from 0.0003 to 0.2, divided in 69 non-linearly spaced intervals.

Simulation-based inference (SBI) is an approach for estimating the probability density of a set of parameters for a given observation in models for which the likelihood is analytically intractable but simulations are possible (Papamakarios and Murray, 2016). Such intractability is present for many models in neuroscience, so SBI has been shown to be a successful approach for performing Bayesian inference in computational neuroscience (Confavreux et al., 2024; Gonçalves et al., 2020). In this work, we followed the algorithms proposed by Gonçalves et al., in which this is achieved by training an artificial neural network to map any simulation result onto a range of possible parameters. To apply this algorithm to our CA3 model, we explored the most relevant parameters of the models, which are the connectivity probability c, ensemble size s, quantal variability σN, threshold g0 and inhibitory weight g1 (because of lack of experimental information). The remaining parameters were fixed to x0 = 1, y0 = 0 and μN = 1 (making CV = σN/μN = σN). In addition, we fixed the total number of synapses to either 2·10^9^ (mouse hippocampal CA3) or 60·10^9^ (human hippocampal CA3), and performed SBI analysis independently for both sizes. The number of neurons was again computed as a function of the connection probability and the number of synapses.

Model implementation was run on a Lenovo ThinkPad L14 Gen 4 with 16 GB of memory, a 13th Gen Intel® Core™ i5-1335U × 12 processor and Mesa Intel® Graphics (RPL-P), over Ubuntu 22.04.3 LTS. Python 3.7.12 and numpy 1.21 were used for all numerical operations, and scipy 1.11 for the normal cumulative distribution. For SBI implementation, torch 1.13 and sbi 0.22 were also used.

### Statistics

All data are reported as mean ± SEM throughout text and legends, or median with interquartile range where specified. Box plots depict data as median (line), 25th and 75th quartile (box) and min/max points (whiskers). Symbols with errors depict mean ± SEM. Proportional datasets were compared using Fisher’s exact test (e.g. for connectivity). As biological datasets rarely exhibit a normal distribution, non-parametric statistical tests for either two-sample (Mann-Whitney test) or multi-sample (Kruskal-Wallis test) data were applied as appropriate, unless specified otherwise. Statistical tests were performed using GraphPad Prism 9, and exact p values were reported throughout the manuscript.

## Acknowledgements

We thank Florian Marr for excellent technical assistance, Christina Altmutter and Julia Flor for technical support, Alois Schlögl for programming, Todor Asenov for development of the transportation box for human brain tissue, Tim Vogels for guidance on simulations, Marcus Huber for mathematical advice, and Eleftheria Kralli-Beller for manuscript editing. This research was supported by the Scientific Services Units (SSUs) of ISTA, and we are particularly grateful for assistance from Christoph Sommer and the Imaging and Optics Facility, Preclinical Facility, Life Science Facility, Miba Machine Shop, and Scientific Computing. We also acknowledge the excellent support of the Medical University of Vienna Department of Neurosurgery staff, Romana Hoeftberger and the Division of Neuropathology and Neurochemistry, and Gregor Kasprian and the Division of Neuroradiology and Musculoskeletal Radiology. The project received funding from the European Research Council (ERC) under the European Union’s Horizon 2020 research and innovation programme (Marie Skłodowska-Curie Actions Individual Fellowship no. 101026635 to J.F.W.), the Austrian Science Fund (FWF; grant PAT 4178023 to P.J.; grant DK W1232 to M.R.T. and J.G.D.) and the Austrian Academy of Sciences (DOC fellowship 26137 to M.R.T.).

## Author Contributions

J.F.W. and P.J. conceived the project, J.F.W., V.V.-B., R.J.M.-M., and M.R.T. performed experiments and analyzed data, K.R. performed epilepsy surgery, A.N.-O. performed modeling, J.G.D. contributed to expansion microscopy analysis, M.T. provided clinical support, J.F.W. and P.J. wrote the paper and acquired funding for the project. All authors jointly revised the paper.

## Supplementary Information

**Supplementary Table 1:** Sample information. Cell Density in *cornu ammonis* (CA) and GC dispersion as reported from routine histological analysis of tissue samples by medical staff. ‘ILAE Scl’ refers to hippocampal sclerosis classifications (see Blümcke et al., 2013).

|  | Class | CA cell density | GC dispersion | ILAE Scl |
| --- | --- | --- | --- | --- |
| 1 | Sclerotic | Almost total nerve cell loss | dispersion | Type 1 |
| 2 | Sclerotic | Subtotal loss of nerve cells | clear cell loss | Type 1 |
| 3 | Non-sclerotic | Regular cell content | none recorded | None |
| 4 | Sclerotic | Almost complete cell loss | slight/noticeable | Type 1 |
| 5 | Non-sclerotic | Regular cell content | possible/slight | None |
| 6 | Non-sclerotic | Regular. CA4: minimal 'extremely discrete' loss | possible/slight | None |
| 7 | Sclerotic | Nerve cell depletion | slight/thinned layer | Type 1 |
| 8 | Non-sclerotic | Regular cell content | negative | None |
| 9 | Sclerotic | Clear nerve cell depletion | not recorded | Type 1 |
| 10 | Non-sclerotic | Regular cell content | negative | None |
| 11 | Sclerotic | Clear nerve cell depletion | slight/thinned layer | Type 1 |
| 12 | Sclerotic | Slight to moderate reduction | negative | Type 2 |
| 13 | Sclerotic | Clear nerve cell loss | dispersed | Type 1 |
| 14 | Non-sclerotic | Regular cell content | negative | None |

**Supplementary Fig. 1.**
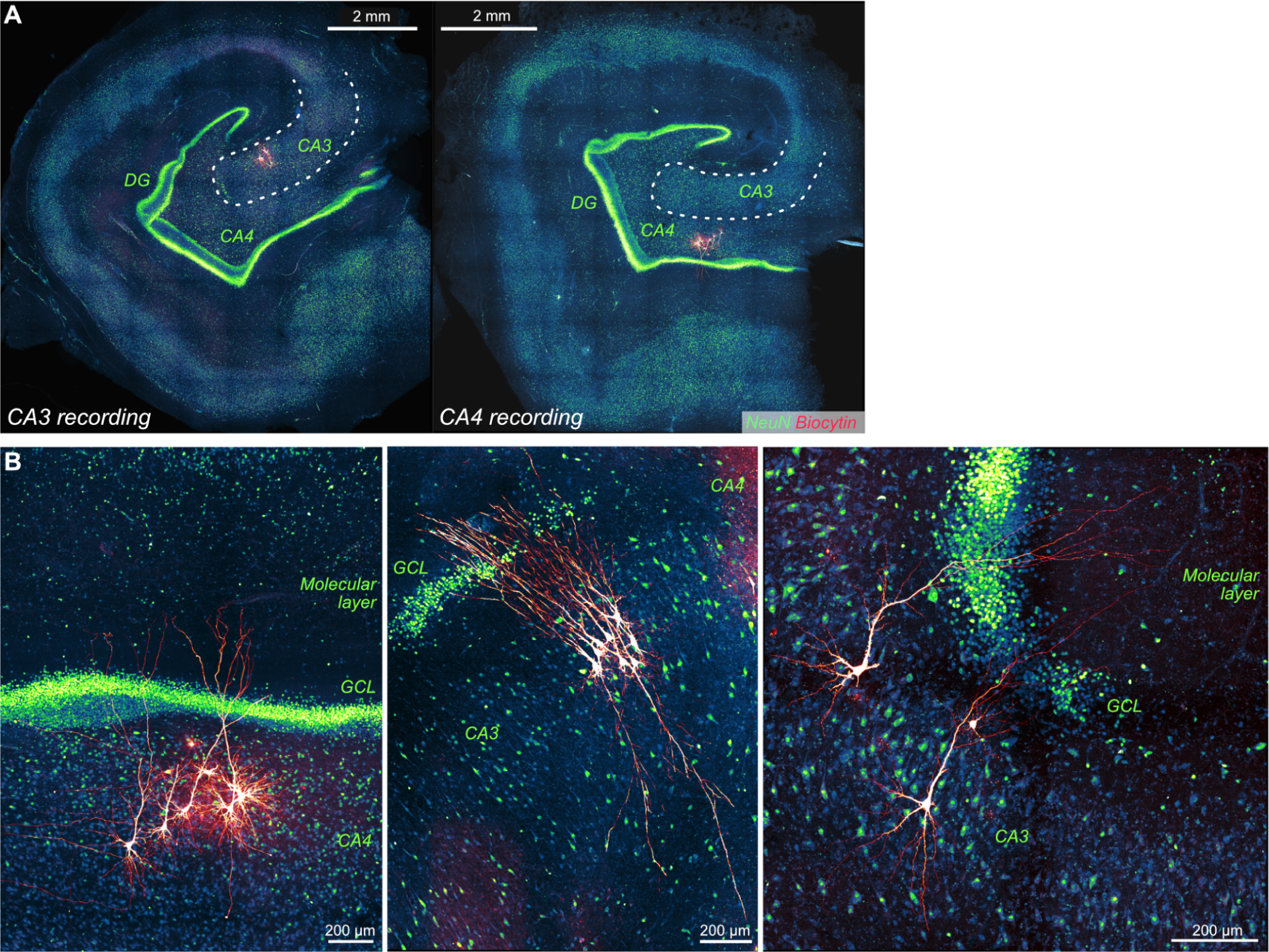
Anatomical features of human hippocampus. **A,** Multicellular recording examples from CA3 (left) and CA4 (right) of the human hippocampus, with CA3 defined as cells within the structured neuronal layer extending out from between the DG granule cell layers (white dashes). **B**, Examples of recorded human CA3/4 PNs showing apical dendrite crossing of the DG granule cell layer (GCL; intracellular biocytin - red, anti-NeuN - blue/green).

**Supplementary Fig. 2.**
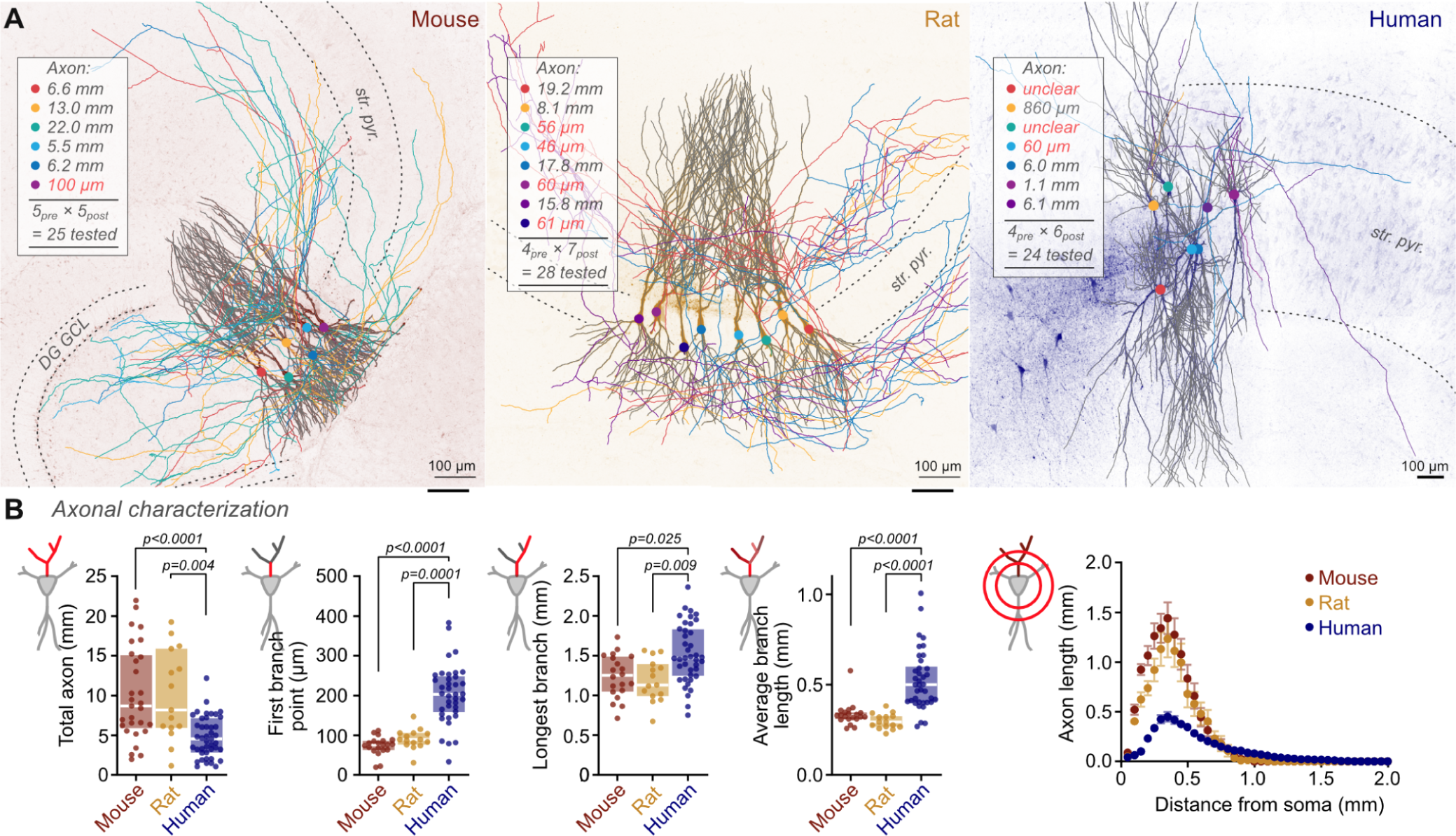
CA3 PN Axonal branch properties across species. **A**, Example multicellular recordings across species to demonstrate axonal cutting analysis. Cells with greater than 0.5 mm of reconstructable axon were included in ‘corrected’ connectivity analysis, which exclude cells with axons exiting the slice before extensive collateralisation, or without visible axons. This criteria will limit slicing-induced underestimation of intercellular connectivity. Maximum intensity projections of streptavidin-AF647 visualised recorded neurons are overlain with reconstructed morphologies. Dendrites are depicted in gray, while individual cell axons/somata are coloured, with quantified length inset. DG granule cell layer (GCL), and CA3 *stratum pyramidale* (str. pyr.) are delineated. **B**, The length of reconstructable axon per neuron is greater in rodent than in human tissue slices (mean ± SEM; mouse: 10.2 ± 1.1 mm; rat: 10.0 ± 1.5 mm; human: 4.8 ± 0.4 mm; Kruskal-Wallis test, p < 0.0001). The reduction in axon length for human CA3 cells results from their less frequent collateralisation. Axonal length to the first branch point is substantially greater in human than rodent CA3 PNs (mouse: 73 ± 5 µm, rat: 90 ± 7 µm, human 203 ± 12 µm; Kruskal-Wallis test, p < 0.0001), and the average axonal length between branch points is also greater in humans than rodents (mouse: 334 ± 14 µm, rat: 297 ± 11 µm, human: 523 ± 27 µm; Kruskal-Wallis test, p < 0.0001). Despite this, the longest axonal branch reconstructable was greatest in human neurons, reflecting the greater size of tissue slice and axonal extent (mouse: 1.24 ± 0.06 mm, rat: 1.29 ± 0.07 mm, human: 1.52 ± 0.06 mm; Kruskal-Wallis test, p = 0.0023). Axonal Sholl analysis demonstrates the reduced axonal extent in human tissue, with a right-shifted distribution, indicative of a broader axonal arborisation. Graphed statistics represent Dunn’s multiple comparisons test throughout. Only cells with > 0.5 mm reconstructable axon were included in **B** and are a subset of recorded cells which are fully reconstructed.

**Supplementary Fig. 3.**
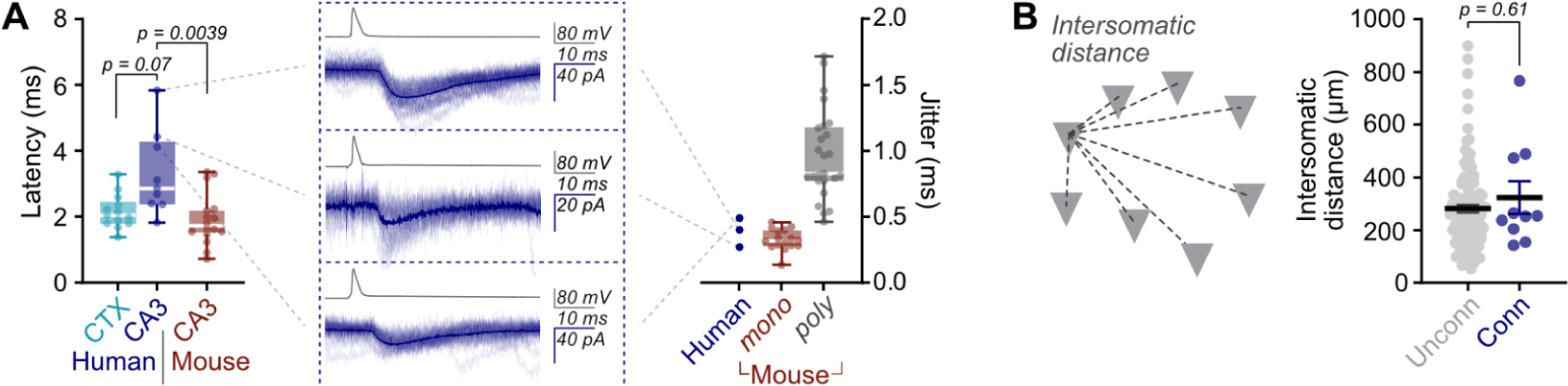
Latency and distance dependence of human CA3 connections. **A**, Latency of human CA3 connections is longer than recorded in mouse CA3 (human cortex: 2.2 ± 0.1 ms, n = 13; human CA3: 3.3 ± 0.4 ms, n = 9; mouse CA3: 1.9 ± 0.2 ms, n = 16; Kruskal-Wallis test, p = 0.0054, Dunn’s multiple comparisons tests depicted). Long latency (> 4 ms) human CA3 connections are likely to be monosynaptic, due to their reliability and low jitter (right). Jitter values (SD of latency) are in line with mouse putative monosynaptic connections and much lower than mouse putative polysynaptic connections (jitter - mouse monosynaptic: 0.32 ± 0.02 ms, n = 16; mouse polysynaptic: 0.97 ± 0.07 ms, n = 25). **B**, No distance dependence to connectivity was observed. Intersomatic distance (3D Euclidean distance between somatic centers) was no different between unconnected and synaptically connected CA3 neurons (unconnected: 281 ± 15 µm, n = 111; connected: 324 ± 62 µm, n = 10, Mann-Whitney test, p = 0.61).

**Supplementary Fig. 4.**
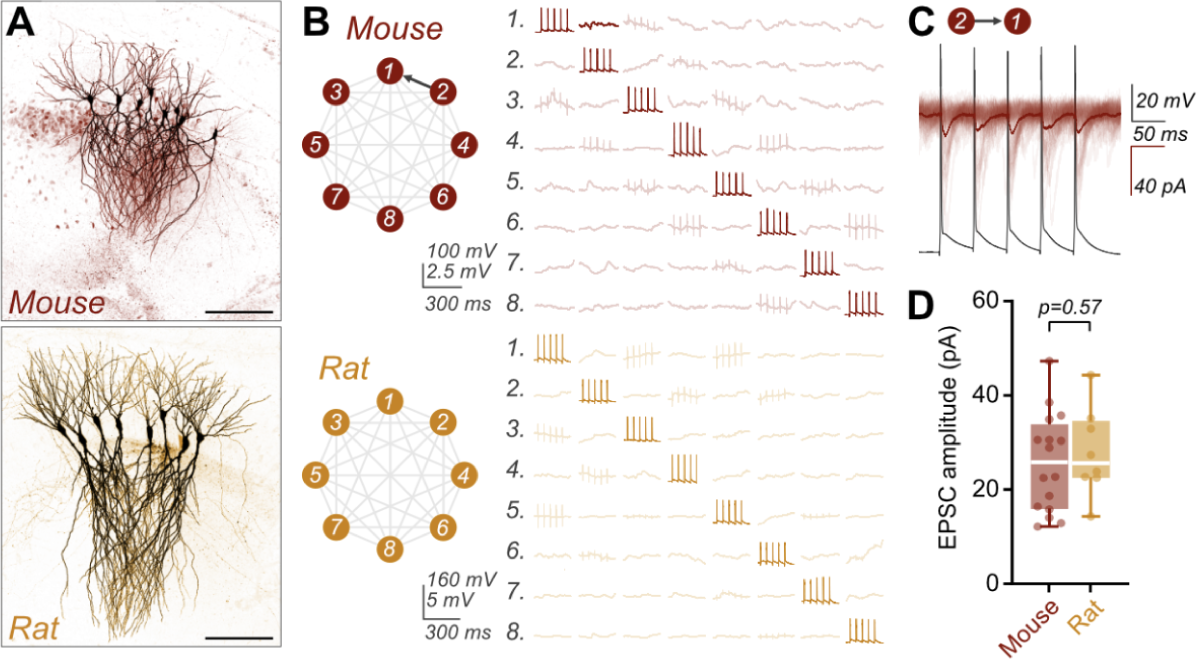
Multicellular recordings from rodent slices. **A**, Example images of octuple patch clamp recorded CA3 cells from mouse (upper) and rat (lower). Scale bar = 200 μm. **B**, Example recordings from mouse and rat CA3 pyramidal neurons. Each recorded neuron is depicted as a row (labeled) of the recording array. Averaged voltage traces in response to presynaptic cell spiking (diagonal) are shown in bold, with, and faded, without, identified monosynaptic connections. Connectivity schemes for these recordings are depicted (left). **C**, Voltage clamp recording of synapse identified in **B**. Individual sweeps (faded) are overlaid with the average trace, with presynaptic cell spiking depicted (gray). **D**, Peak amplitudes of unitary EPSCs are similar between mouse and rat connections (mouse: 25.8 ± 2.6 pA, n = 16; rat: 27.9 ± 3.3 pA, n = 8; Mann-Whitney test, p = 0.57).

**Supplementary Fig. 5.**
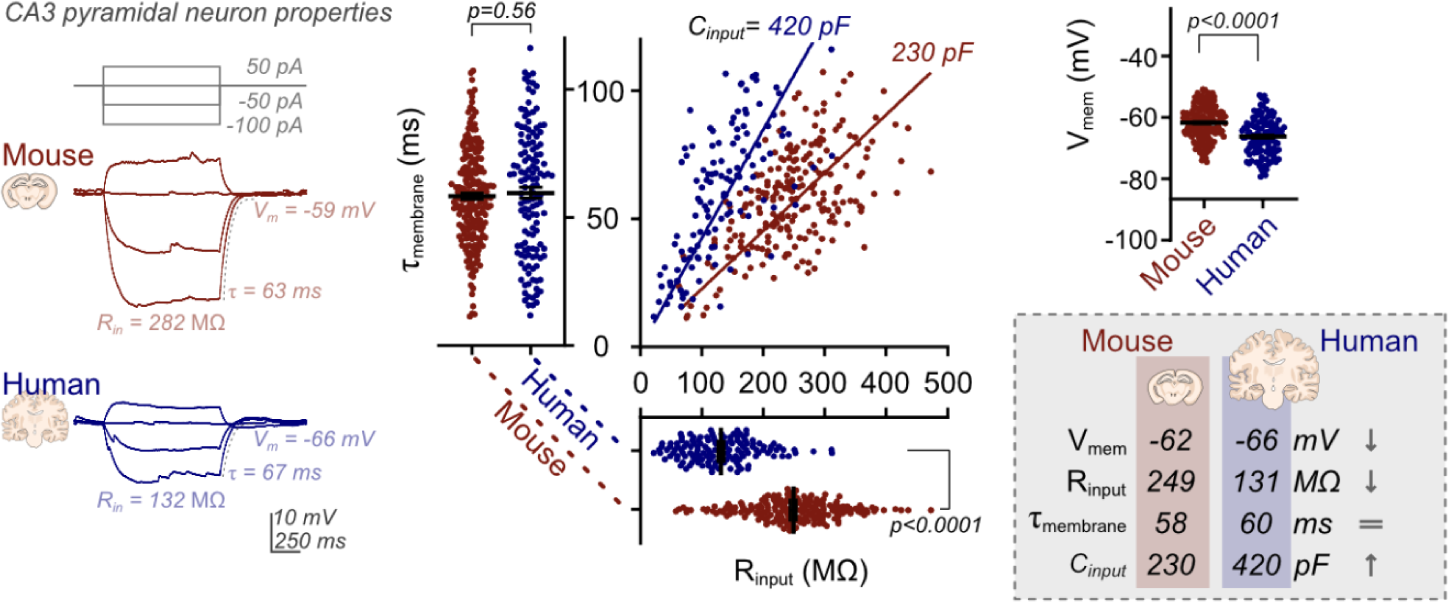
CA3 PN passive membrane properties across species. Human CA3 PNs have lower input resistance (mouse: 249 ± 5 MΩ, n = 232; human: 131 ± 5 MΩ, n = 135; Mann-Whitney test, p < 0.0001) but similar membrane time constant (mouse: 58.3 ± 1.2 ms, n = 229; human: 59.6 ± 2.2 ms, n = 133; Mann-Whitney test, p = 0.56) to mouse CA3 PNs, which correspond to an increased cell capacitance (linear fit of input resistance vs. membrane time constant scatter - human: 420 pF, mouse 230 pF). The resting membrane potential of human CA3 PNs is subtly lower than that of mice (mouse: −61.8 ± 0.4 mV, n = 232; human: −66.2 ± 0.5 mV, n = 135; Mann-Whitney test, p < 0.0001).

**Supplementary Fig. 6.**
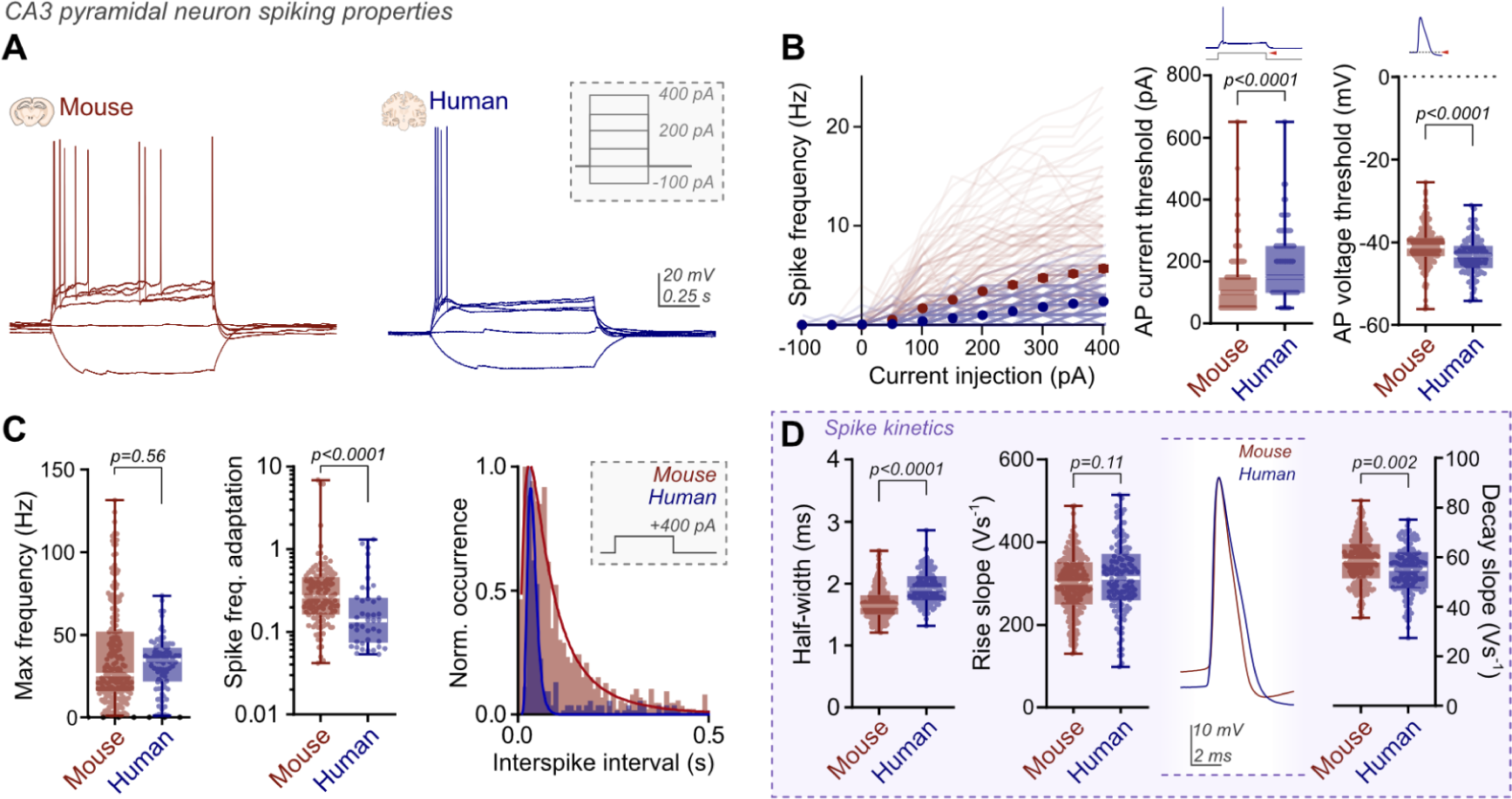
CA3 PN active membrane properties across species. **A**, Example traces of mouse and human action potential responses to current injection (inset). **B**, Mouse neurons give greater spiking (across 1-s stimulus, symbols - mean ± SEM, lines - individual cells), with lower current threshold for spike generation (rheobase / minimum current for spiking - mouse: 125 ± 5 pA, n = 232; human: 197 ± 9 pA, n = 134, Mann-Whitney test, p < 0.0001). Action potential threshold is subtly lower in human PNs (mouse: = −40.9 ± 0.3 mV, n = 232; human: −43.4 ± 0.4 mV, n = 134, Mann-Whitney test, p < 0.0001). **C**, CA3 neurons exhibit maximal spiking at current onset, which is more pronounced in human neurons, as observed through multiple parameters. Maximum firing rate at 400 pA injection have similar mean values between species (mouse: 37 ± 2 Hz, n = 212; human: 32 ± 2 Hz, n = 89), however with heterogeneous distributions. The majority of mouse neurons fire at lower frequency than human neurons. Spike frequency adaptation is far greater in human neurons (mouse: 0.47 ± 0.08, n = 128; humans: 0.25 ± 0.05, n = 42, Mann-Whitney test, p < 0.0001). The distribution of inter-spike intervals across all cells (400 pA injection) is much narrower for human neurons, demonstrating more precise spike timing. **D**, The half-width of human spikes is subtly longer than those of mice (mouse: 1.68 ± 0.02 ms, n = 232; human: 1.94 ± 0.02 ms, n = 134; Mann-Whitney test, p < 0.0001), with more change in action potential maximal decay slope (mouse: 58.6 ± 0.6 V s^−1^, n = 232; human: 54.8 ± 0.8 V s^−1^, n = 134; Mann-Whitney test, p = 0.002) than rise slope (mouse: 298 ± 5 V s^−1^, n = 231; human: 314 ± 8 V s^−1^, n = 134; Mann-Whitney test, p = 0.11).

**Supplementary Fig. 7.**
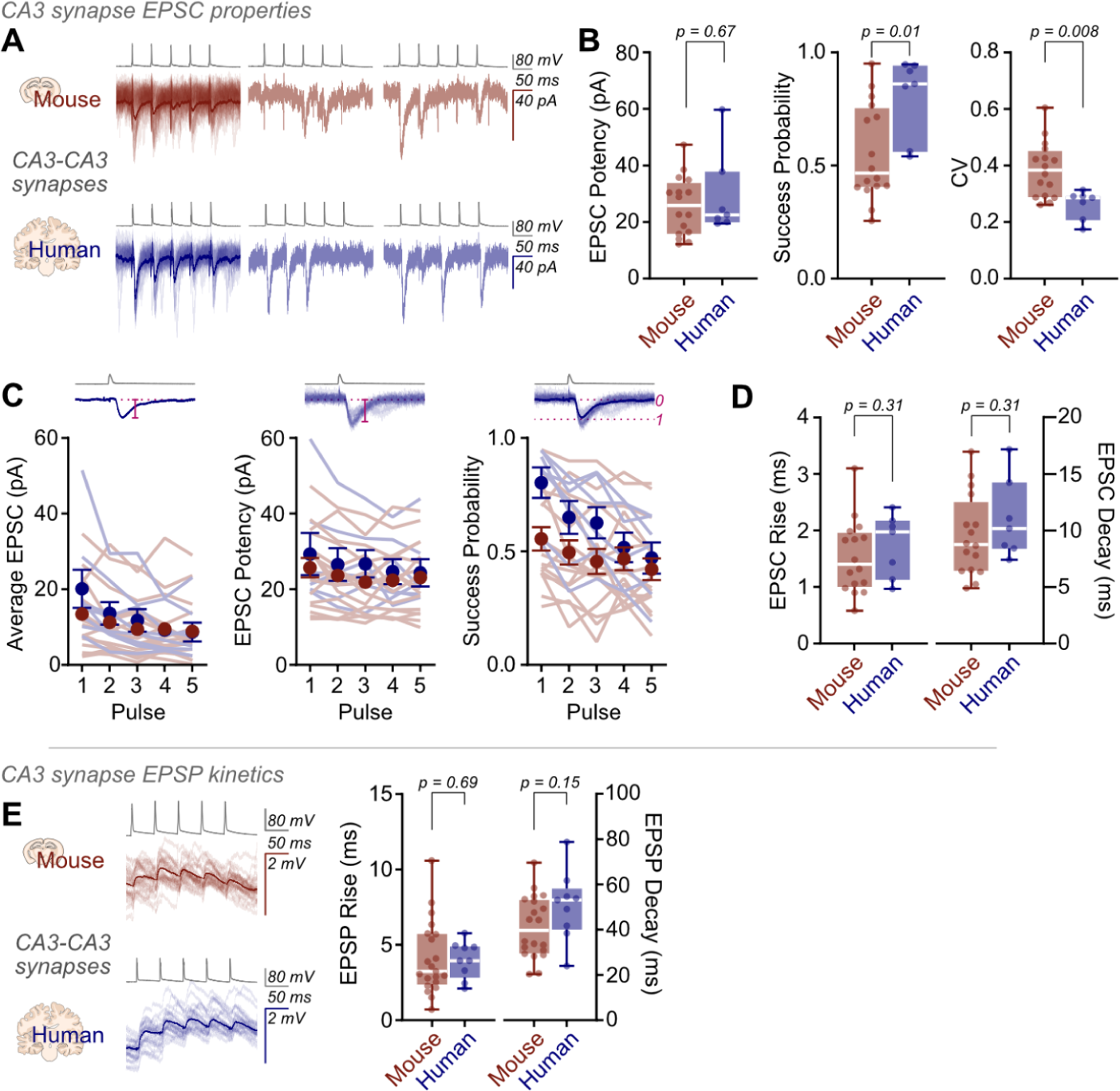
Increased synaptic reliability in the human brain. **A**, Example traces of unitary EPSCs from mouse (upper) and human (lower) CA3 PN pairs. Average traces (bold) are depicted over individual traces (left), with example sweeps presented (right). **B**, Average EPSC potency was unchanged between species (mouse: 25.8 ± 2.6 pA, n = 16; human: 29.4 ± 5.6 pA, n = 7; Mann-Whitney test, p = 0.61), while success probability was significantly higher (mouse: 0.56 ± 0.05, n = 16; human: 0.80 ± 0.07, n = 7; Mann-Whitney test, p = 0.010) and coefficient-of-variation (CV) of success amplitudes significantly lower in human synapses (mouse: 0.39 ± 0.02, n = 16; human: 0.26 ± 0.02, n = 7; Mann-Whitney test, p = 0.008). **C**, Individual (lines) and average (circles) responses to 20-Hz train stimulation. Average EPSCs (left), are produced from EPSCs of a similar potency throughout trains and between species (center), with different success probabilities (right). **D**, EPSC kinetics appeared unchanged between mouse and human CA3 synapses (EPSC rise time (20–80%) - mouse: 1.54 ± 0.16 ms, n = 16; human: 1.74 ± 0.21, n = 7; Mann-Whitney test, p = 0.31. EPSC decay time constant - mouse: 9.55 ± 0.90 ms, n = 16; human: 11.02 ± 1.33, n = 7; Mann-Whitney test, p = 0.31). **E,** EPSPs (20-Hz train stimulation traces) of unitary CA3 synapses have similar kinetics between mice and humans (EPSP 20–80% rise time - mouse: 4.1 ± 0.5 ms, n = 21; human: 4.0 ± 0.4 ms, n = 9; Mann-Whitney test p = 0.69. EPSP decay time constant - mouse: 41.1 ± 3.1 ms, n = 20; human: 50.6 ± 5.1, n = 9; Mann-Whitney test, p = 0.15).

**Supplementary Fig. 8.**
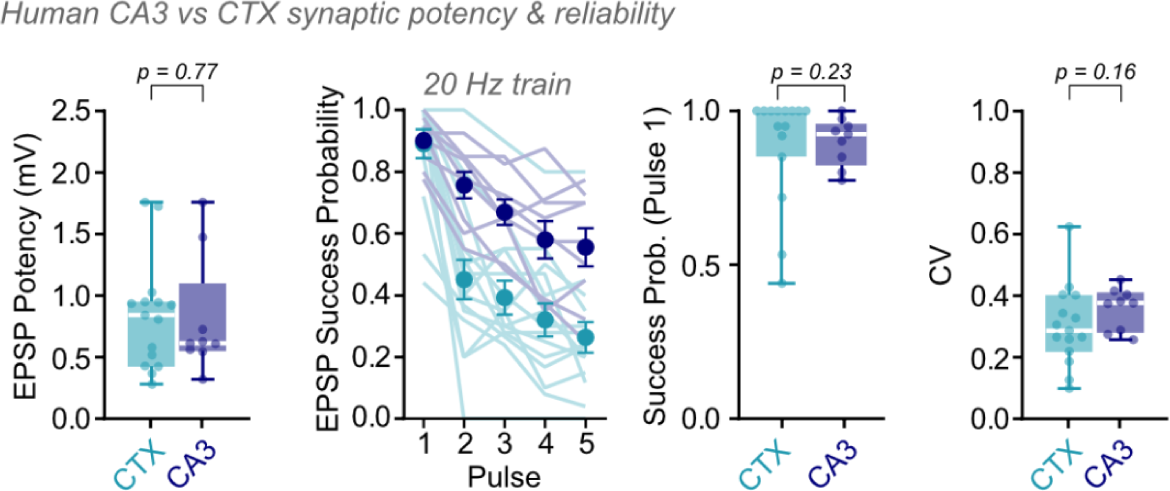
Human CA3 and neocortical synapses showed not only similar potency (CTX: 0.84 ± 0.11 mV, n = 15; CA3: 0.80 ± 0.16, n = 9, Mann-Whitney test, p = 0.77), but also similarly high success probability for the first pulse of 20-Hz trains (CTX: 0.89 ± 0.05, n = 15; CA3: 0.90 ± 0.03, n = 9; Mann-Whitney test, p = 0.23) and similarly low CV (CTX: 0.30 ± 0.03, n = 15; CA3: 0.36 ± 0.02, n = 9; Mann-Whitney test, p = 0.16), suggesting that synaptic reliability is a feature of the human brain.

**Supplementary Fig. 9.**
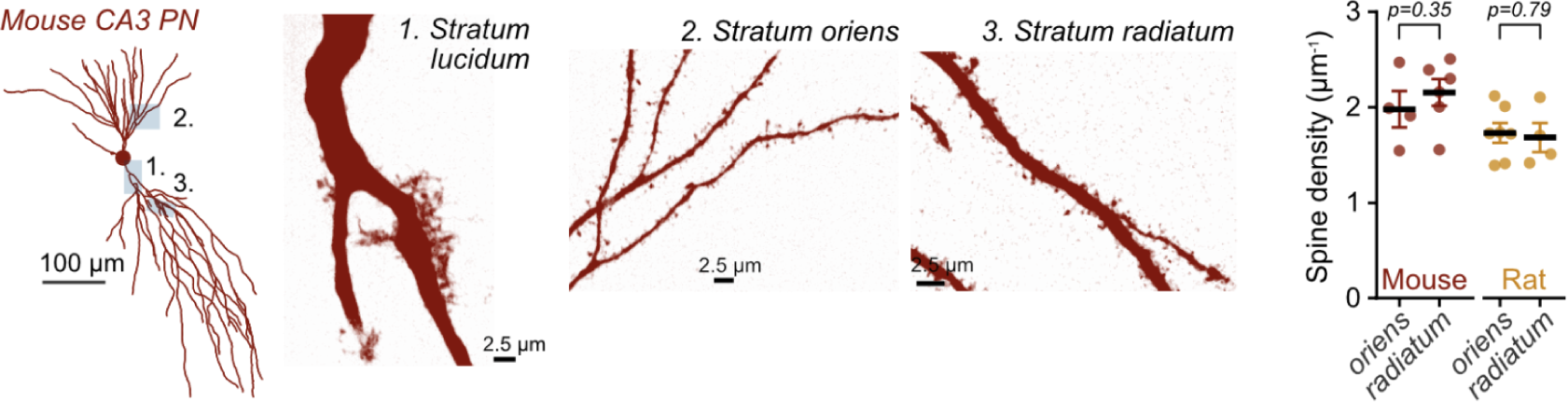
Spine density is similar between CA3 layers. Reconstructed mouse neuron (left, skeleton), with labeled areas imaged by 4x expansion microscopy (center). Image scale bars depict pre-expansion sizes. Measured spine density (per μm dendrite) shows no difference between dendrites in *stratum oriens* or *radiatum* in either mouse or rat samples (mean ± SEM, mouse *stratum oriens*: 1.98 ± 0.19 µm^−1^, n = 4; *stratum radiatum*: 2.16 ± 0.14 µm^−1^, n = 6; Mann-Whitney test, p = 0.35. Rat *stratum oriens*: 1.73 ± 0.10 µm^−1^, n = 7; *stratum radiatum*: 1.68 ± 0.15 µm^−1^, n = 4; Mann-Whitney test, p = 0.79).

**Supplementary Fig. 10.**
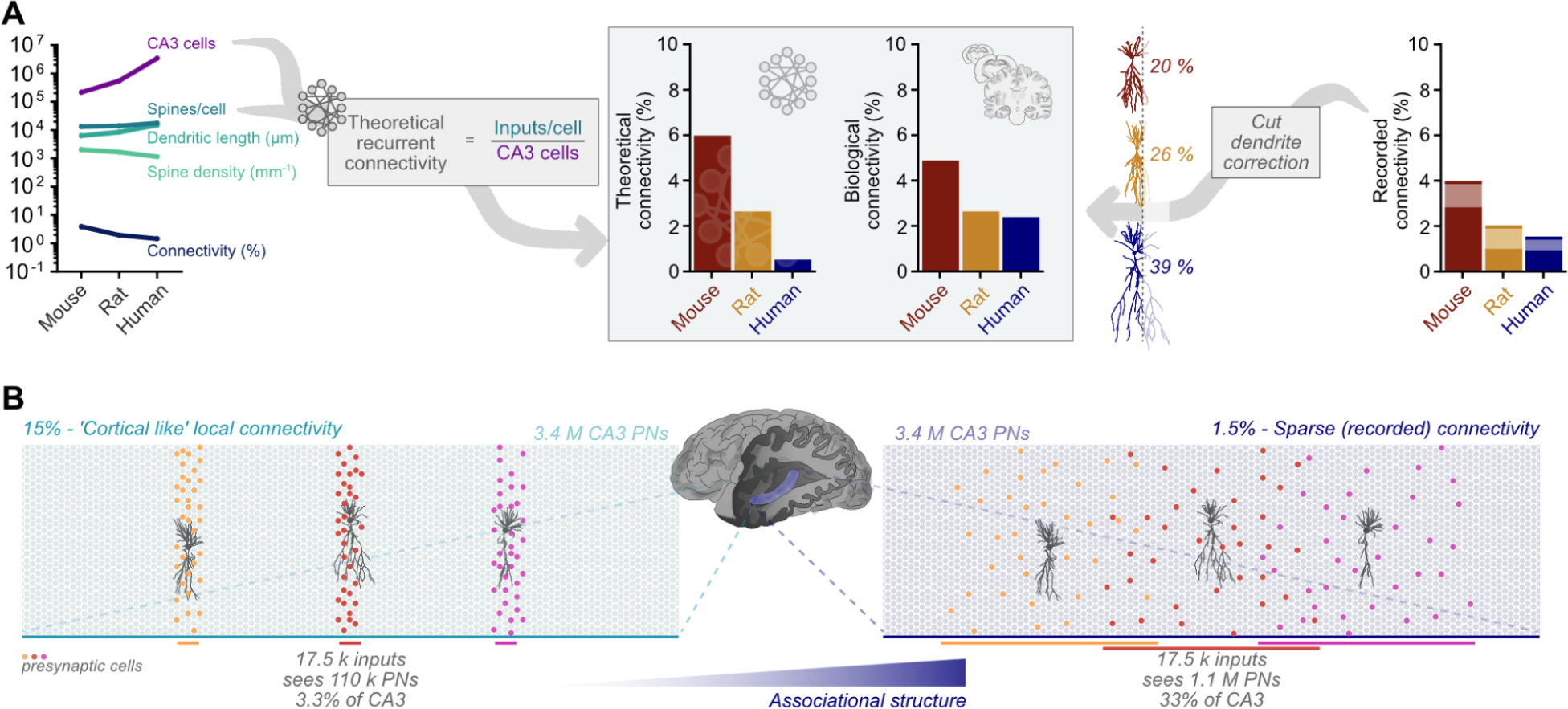
Theoretical analysis of recurrent collateral synapses predicts sparse connectivity. **A**, Theoretical connectivity levels for a broad recurrent network can be calculated anatomically from the number of network nodes (CA3 neurons) and inputs per node (spines/cell), which predicts mouse, rat, and human CA3 connectivity at 5.99%, 2.65%, and 0.52%, respectively. These values are comparable to the estimated connectivity from recordings, indicating that CA3 approximates a broad recurrent network across species (connectivity after correction for both axon cutting (as previous), and percentage of recorded cell dendrites lost by slicing gives ‘biological connectivities’ - mouse: 4.89%, rat: 2.65%, human: 2.41%). **B**, Sparse CA3 connectivity forms a broad associational network. Dense, neocortical levels of local connectivity (left) would imply that individual CA3 PNs receive all 17,500 inputs from just 3.3% of the total CA3 volume. Sparse local connectivity, as recorded (right), allows for broader, long-range influence on individual CA3 PNs, forming a broad associational network. Colors represent putative presynaptic cells to each CA3 PN, constrained by dense or sparse connection density. Human brain schematic depicting hippocampus (blue), based on Gray’s Anatomy of the Human Body (1918).

